# Early micro and nanoscopic responses of microglia to blood-brain barrier modulation by transcranial-focused ultrasound

**DOI:** 10.1101/2025.09.17.675183

**Authors:** Elisa Gonçalves de Andrade, Jared VanderZwaag, Rikke Hahn Kofoed, Micaël Carrier, Katherine Picard, Keelin Henderson Pekarik, Fernando González Ibáñez, Mohammadparsa Khakpour, Kullervo Hynynen, Isabelle Aubert, Marie-Ève Tremblay

**Author notes:** Co-correspondence: Marie-Ève Tremblay, Isabelle Aubert.

## Abstract

Modulation of the blood-brain barrier (BBB) using transcranial-focused ultrasound (FUS) has rapidly progressed to clinical trials. In combination with phospholipid microspheres, also known as microbubbles, administered in the bloodstream, ultrasound energy is guided by magnetic resonance imaging (MRI) to target specific brain regions with millimetric precision. At the targeted area, the interaction between FUS and microbubbles increases local BBB permeability for 4 to 6 hours, with an ensuing inflammation that resolves within days to weeks. Microglia, as the resident immune cells of the brain, are triggered by FUS-BBB modulation, although the time course of this response is unclear. Thus, the goal of this study was to characterize the early cellular (i.e., density, distribution, and morphology) and subcellular (i.e., ultrastructure) changes in microglial activities following FUS-BBB modulation.

**Methods:** We targeted the hippocampi of adult mice with FUS, in the presence of intravenous microbubbles and guided by MRI, and performed analyses 1 hour and 24 hours after FUS-BBB modulation. Microglia were investigated at the population, cellular and subcellular levels, where hippocampal BBB permeability was identified by the entry of endogenous immunoglobulin (Ig)G in the parenchyma. Respective outcome measures included i) the density and distribution of ionized calcium binding adaptor molecule-positive (Iba)1-positive (+) cells; ii) the morphology of the soma and processes of Iba1+ cells; and iii) the quantification of microglial organelles (e.g., phagosomes) and contacts with blood vessels and synapses using chip mapping scanning electron microscopy.

**Results:** No significant changes in baseline density and distribution of microglia were found in IgG-positive hippocampal areas at 1 hour and 24 hours after FUS-BBB modulation. By contrast, FUS-BBB modulation was associated with more elongated microglial cell bodies at both time points. The relative distribution of morphologies at 1 hour shifted toward compact shapes with stubby processes, whereas at 24 hours, shapes were bigger, with fewer processes. At the nanoscale, microglia maintained their interactions with blood vessel elements, except vessels most affected by swollen endfeet, which occurred regardless of treatment. In the parenchyma, 24 hours after FUS-BBB modulation, microglia reduced the frequency of contacts with pre-synaptic elements and extracellular space pockets, while showing features of increased metabolic demand and reduced lysosomal activity.

**Conclusion:** At 1 hour and 24 hours after FUS-BBB modulation, traits of microglial surveillance activity were largely maintained, with shifts in the shape of a subset of cells, which adopted a morphology associated with injury shielding. FUS-BBB modulation also appears to temporarily modify the digestive, but not the phagocytic activity, of microglia and to reduce pre-synaptic remodeling early after treatment.

## Introduction

Microglia are the resident immune cells of the central nervous system (CNS), which play a key role in neuronal, vascular and inflammatory pathology across brain diseases. [1,2]. Changes in the activity of microglia, commonly known as ‘reactivity’, can impact the CNS through altered brain surveillance, synaptic remodeling, and blood-brain barrier (BBB) maintenance [3,4]. Microglia coexist in a spectrum of states presenting many levels of complexity, pertaining to molecular expression, ultrastructure, morphology, distribution and density [3–5]. These levels of complexity together determine microglial functions, making their detailed characterization necessary for identifying strategies to retaining beneficial physiological roles.

Focused ultrasound (FUS) combined with intravenously injected microbubbles and guided by magnetic resonance imaging (MRI) is a cutting-edge technology that modulates the BBB, and improves numerous neurological conditions. The BBB is a dynamic border checkpoint of the CNS, regulating the flux of molecules and cells into the brain parenchyma [8–10]. FUS-BBB modulation transiently increases BBB permeability, improving the entry of therapeutic drugs into the CNS of rodents [11], non-human primates [12] and humans undergoing clinical trials [11]. In rodent models of neurodegenerative diseases, FUS-BBB modulation has been associated with beneficial microglial responses days after treatment, possibly facilitating the removal of deleterious molecules by modulating their surveillance and phagocytic activities [13]. The opposite may be true in the healthy brain, where reduced microglial proximity and cellular processes were found in the targeted compared to non-targeted hippocampi of mice 7 days after FUS-BBB modulation [14]. Evidence of a similar effect early after FUS-BBB modulation is lacking.

Moreover, transient shifts in microglial morphologies associated with reduced homeostatic parenchymal surveillance may be a contentious point of FUS-BBB modulation [15].

During acute inflammatory processes microglia shift at the nanoscale level and present with increased ultrastructural markers of cellular stress [16], including dilated endoplasmic reticulum (ER)/Golgi cisternae and enlarged mitochondria [17,18]. Microglia showing ultrastructural markers of stress in neurodegenerative mouse models interact less with the vasculature [19] and healthy microglia seem to be required for timely closure of the BBB following FUS [20]. Thus, during the window of FUS-induced BBB permeability, typically lasting 4 to 6 hours [9,21], microglia under cellular stress could alter their physiological functions and impact the resolution of BBB properties.

To provide insights into the early changes in microglial features, including surveillance and phagocytosis, following FUS-BBB modulation in adult mice, we targeted a memory and learning hub, the hippocampus, and collected brains at 1 hour and 24 hours post-FUS-BBB modulation. Sections were stained for ionized calcium-binding adaptor molecule (Iba)1, a microglia/macrophage marker, and immunoglobulin (Ig)G, which we used as an indicator of increased BBB permeability [13,22]. Using brightfield microscopy, we measured the density, distribution, and morphology of Iba1+ cells. With chip mapping scanning electron microscopy (SEM), we assessed microglial intracellular organelles (e.g., phagosomes) and contacts with the BBB and neuronal elements. Our findings support that FUS-BBB modulation does not modify the density and distribution of hippocampal microglia in the first 24 hours of treatment. During this time, microglial cell bodies were enlarged, and a subset of cells shifted to larger shapes, with fewer processes. These modifications were accompanied by more frequent interactions with vessels showing swollen astrocytic endfeet, but the opposite effect on contacts with pre-synaptic elements and extracellular space pockets. In addition, as features of metabolic demand increased (i.e., elevated mitochondrial alterations), microglia appeared to reduce their lysosomal efficiency. Therefore, our results suggest that FUS-BBB modulation does not disrupt microglial surveillance and phagocytosis and that a subset of cells adapts to BBB permeability, with potential effects on synaptic plasticity and other physiological roles.

## Results

### FUS-BBB modulation increased hippocampal BBB permeability

Microglia work in unison with the other cells of the neurovascular unit, aiding in blood vessel dilation and constriction, as well as BBB monitoring and repair, through the exchange of extracellular matrix molecules and direct cell-to-cell contacts [15]. During increased BBB permeability following FUS, these microglial roles could change in response to the influx of blood-derived signals into the brain, such as IgGs [15]. To provide insights into these possible changes in microglial activities, the hippocampi of adult mice (n = 6 animals) were targeted ipsilaterally with FUS.

The corresponding contralateral region was used as a control. Immediately after FUS-BBB modulation, gadolinium-enhanced T1-weighted MRI was acquired to confirm increased BBB permeability in the targeted hippocampi (Figure 1A). MR images were used to quantify parenchymal gadolinium leakage (Figure 1A, dashed circle), where we detected a significantly increased average voxel intensity in the ipsilateral compared to the contralateral hippocampi (Figure 1B *p =* 0.0449, t = 2.661, Table S1).

**Figure 1.**
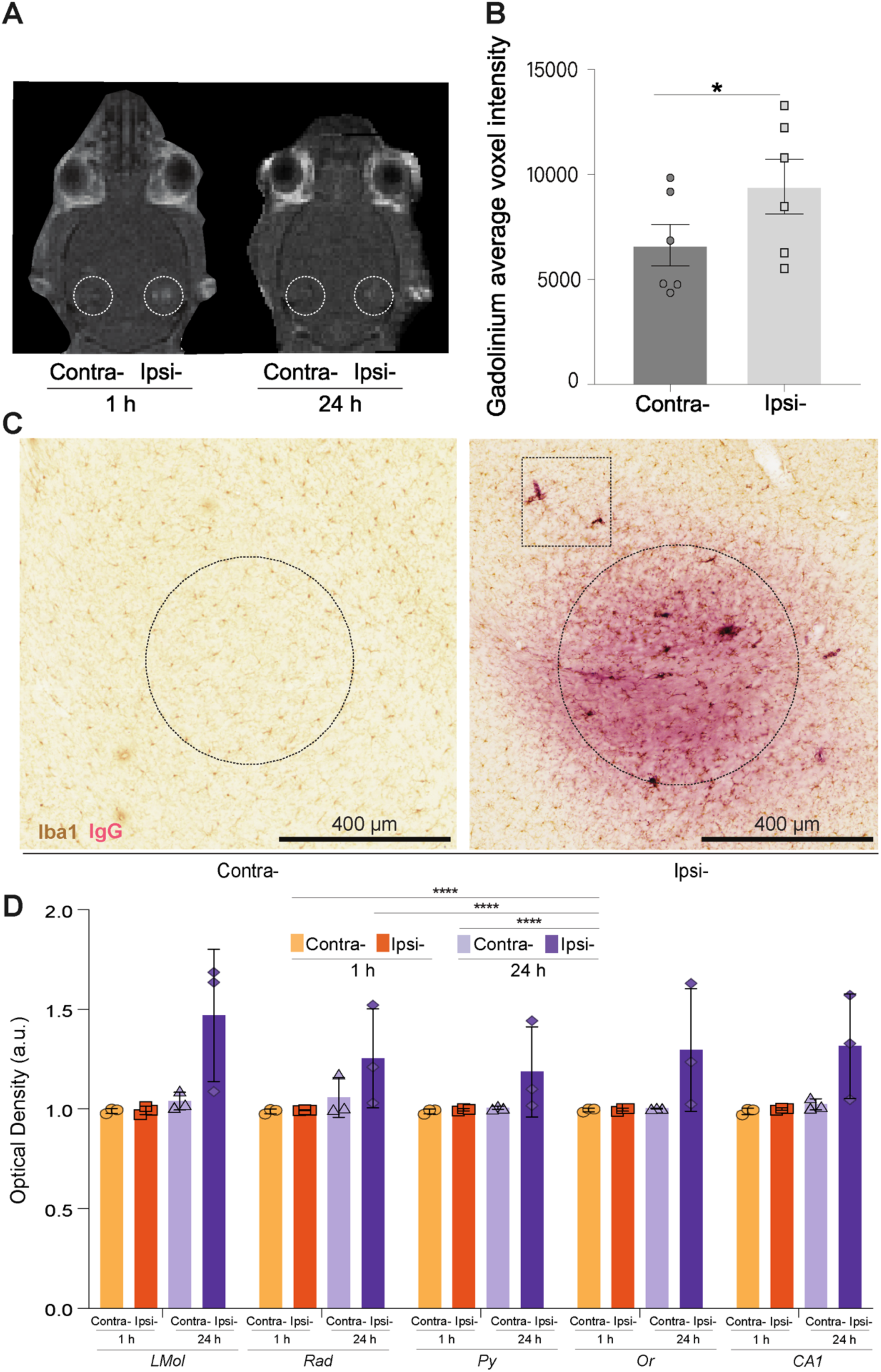
FUS-BBB modulation increased hippocampal BBB permeability. **A.** T1-weighted images confirmed increased blood-brain barrier (BBB) permeability indicated by gadolinium detection (dashed circle) in the ipsilateral (ipsi-) hippocampi parenchyma at 1 hour (h) and 24 h after FUS-BBB modulation. **B.** When quantified, the average voxel intensity of the gadolinium signal was significantly increased in the ipsilateral compared to the contralateral (contra-) hippocampi. The bar graph shows the mean, standard error of the mean, and individual data points (n = 3 animals/hemisphere/time point), analyzed with a paired two-tailed Student’s *t*-test. **C.** Representative images of ionized calcium-binding adapter molecule (Iba)1, a marker of microglia/macrophages, and immunoglobulin (Ig)G immunoperoxidase staining imaged in the *cornu ammonis (CA)1* at 40x with a brightfield microscope. IgG was only detected in the ipsilateral parenchyma and was often circular (dashed circle), indicative of the area targeted by the FUS-BBB modulation. Scale bars 400 μm. **D.** Optical density was significantly increased across time (main effect of Time not shown) with higher values at 24 h compared to 1 h and baseline, irrespective of layer (*LMol*, *Rad, Py*, *Or*, *CA1*). The bar graphs show the mean, standard error of the mean, individual data points (n = 3 animals/hemisphere/time point), analyzed with a mixed effects 2-way ANOVA and Šídák’s multiple comparison tests. * *p <* 0.05, ** *p <* 0.01, *** *p <* 0.001, **** *p* < 0.0001. a.u.: arbitrary unit, *Or*: *stratum oriens*, *Py*: *stratum pyramidale* and *Rad*: *stratum radiatum*.

Gadolinium parenchymal leakage has long been used to infer real-time increases in BBB permeability [6]; however, to gain insight into the extent of permeability *post-mortem*, we probed the entry of endogenous IgGs from the blood to the brain via immunoperoxidase staining and brightfield microscopy [13,22]. We exclusively observed parenchymal IgG staining in the ipsilateral hippocampi (Figure 1C), particularly in the *cornu ammonis (CA)1*. As the *CA1* is organized in four strata with distinct neuronal, microglial and vasculature compositions, we examined the parenchymal distribution of endogenous IgG following FUS-BBB modulation in these layers (Figure 1D). Medially to laterally, these layers are distributed as: *stratum lacunosum*-*moleculare* (*LMol)*, *stratum radiatum (Rad)*, *stratum pyramidale (Py)* and *stratum oriens (Or)* [23]. As this is an exploratory study, we performed all possible pairwise comparisons, outlined in the supplementary tables. We report in the text, however, only the differences between ipsilateral and contralateral hippocampi across the two time points. We identified increased optical density across time and in the 24 h ipsilateral hemisphere compared to the 1 h ipsilateral, the 1 h and 24 h contralateral hemispheres post-FUS-BBB modulation (Figure 1D, main effect of Time F(3, 30) = 17.49 and *p =* 0.0001, post-hoc contralateral 1 h *vs* ipsilateral 24 h *p =* <0.0001, post-hoc ipsilateral 1 h *vs* ipsilateral 24 h *p =* <0.0001, post-hoc contralateral 24 h *vs* ipsilateral 24 h *p =* <0.0001, Table S1). There was no significant difference across layers (Table S1). Our results overall suggest that hippocampal BBB permeability increased rapidly after FUS-BBB modulation (Figure 1A, B), allowing circulating endogenous IgG to enter and accumulate in the brain parenchyma for at least 24 h.

### Microglial density and distribution were maintained

Microglia dynamically rearrange their distribution, a process which can be influenced by stressors, such as laser injury in mice [24], and by FUS-BBB modulation as observed after 1 and 3 days [20] or 7 weeks [14,25]. To investigate how early these changes appear, we quantified the number and proximity of microglia in individual *CA1* layers of the ipsilateral and contralateral hippocampi post-FUS-BBB modulation (Figure 2A–D). Specifically, we probed Iba1+ cell density, cluster cell density (<12 µm apart), nearest neighbour distance (NND, nearest cell to every other cell), and spacing index (NDD and density compiled). In the different layers examined, there was no significant difference between the contralateral and ipsilateral regions in any of these metrics (Figure 2E-F, Table S2). In the pooled *CA1*, we identified that time modulated the NND (Figure 2G, main effect of Time p = 0.0464 and f = 5.5390), indicating the distance between cells changed over the 24-hour period after FUS-BBB modulation_._

**Figure 2.**
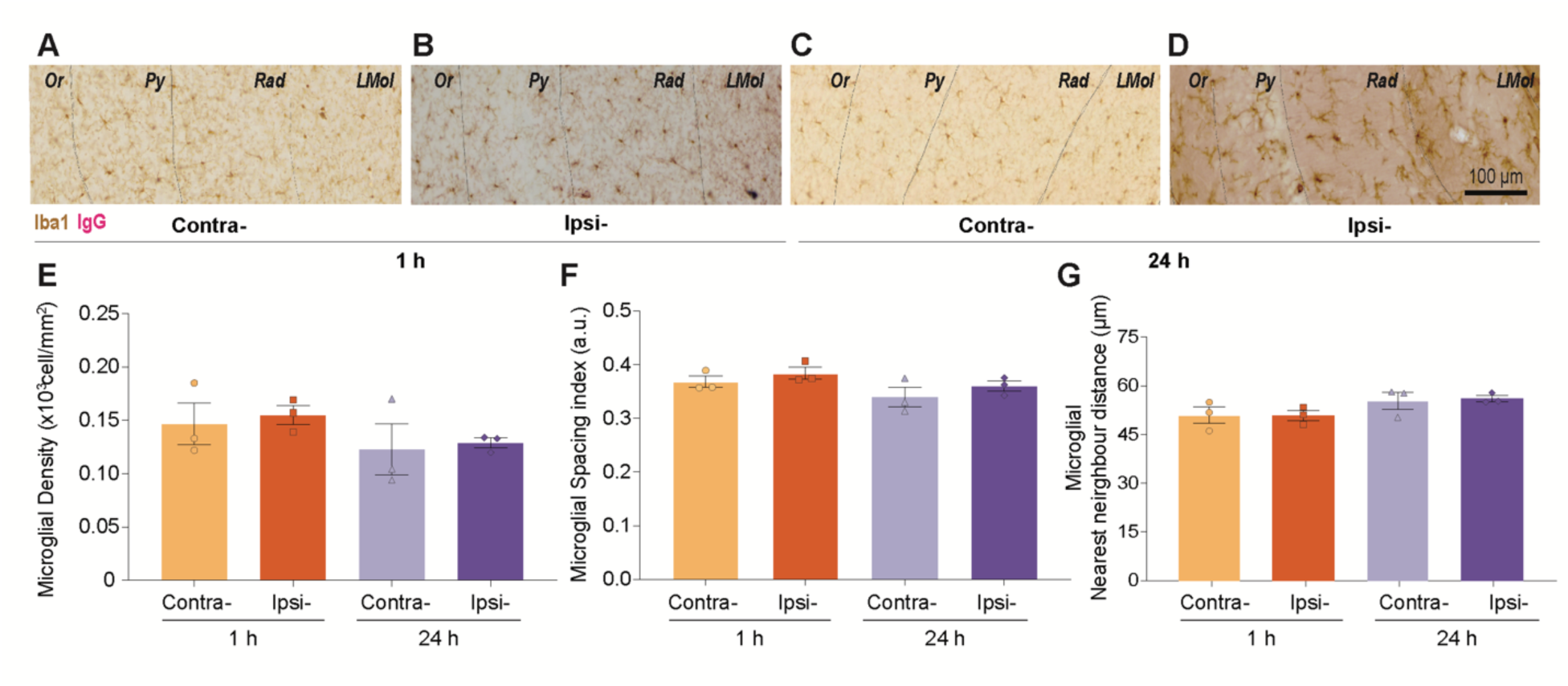
Within one day of FUS-BBB modulation, there were no microglial density or distribution changes. **A-D.** Representative brightfield microscopy images taken at 40x illustrate ionized calcium-binding adapter molecule (Iba)1 positive (+) cell and immunoglobulin (Ig)G + parenchymal distribution along the examined *cornu ammonis (CA)1* strata. Images were obtained in ipsilateral (ipsi-) and contralateral (contra-) hemispheres at 1 hour (h) and 24 h following double immunoperoxidase staining. Scale bar 100 μm. **E-G.** In the *CA1*, the density and spacing index were not altered despite a main effect of Time (not shown) in changing the nearest neighbour distance during the 24 h after FUS-BBB modulation (**G**). The bar graphs show the mean, standard error of the mean, and individual data points (n = 3 animals/hemisphere/time point). Statistical significance was assessed by a mixed-effects 2-way ANOVA with Šídák’s multiple comparison tests. a.u.: arbitrary unit.

Changes in microglial distribution, including NND, may occur in proportion to the extravasation of blood molecules into the CNS [15]. To evaluate this possibility, we examined the Spearman r correlation coefficients of the IgG staining optical density and microglial properties (i.e., cell and cluster density, NND, and spacing index) in the ipsilateral *CA1* at both 1 h and 24 h after FUS-BBB modulation. We observed a significant negative correlation between IgG optical density and microglial density (Figure S1A, r = -0.94 and *p* = 0.02, Table S3) and a positive correlation between IgG optical density and microglial NND (Figure S1A, r = 0.89 and *p* = 0.03, Table S3) at both time points. While our results suggest an inverse relationship between the IgG leakage and microglial proximity, a longitudinal analysis of microglia and blood vessels during the hours and days following FUS-BBB modulation is warranted to assess microglia-vessel leakage relationships [26,27].

### Microglial morphology diversified

Microglia can increase their cell body size in response to elevated gene transcription and protein translation [15]. In rodents, FUS-BBB modulation is associated with an early increase in cytokine expression [28–30] and an elevated soma area in hippocampal microglia 7 days after FUS-BBB [14]. We examined whether shifts in cell body morphology also occur earlier, at 1 h and 24 h after FUS-BBB, in the *CA1 LMol.* Compared to the remaining *CA1* layers, the *LMol* holds large vessels, such as the internal transverse hippocampal vein and external transverse hippocampal artery [31], which may be more heavily impacted by FUS-BBB modulation with microbubbles [32]. We randomly selected and manually traced *LMol* Iba1+ cell bodies (Figure 3A-D), inspecting a variety of shape descriptors. We observed an increase in soma perimeter in the ipsilateral compared to the contralateral *LMol* at 24 h (Figure 3E, main effect of Hemisphere *p =* 0.0171 and F = 15.4300, interaction effect of Hemisphere x Time *p =* 0.0474 and F = 8.0070, post-hoc contralateral vs ipsilateral at 24 h *p =* 0.0175 and *t* = 4.7780, Table S4). This was a specific effect, as the average soma area (Figure 3F, Table S4), aspect ratio (circular *vs* elongated, Figure 3G, Table S4), solidity (porous *vs* circular, Figure 3H, Table S4), roundness (Figure 3I, Table S4), and circularity (Figure 3J, Table S4) were not significantly altered ipsilaterally at 1 h and 24 h after FUS-BBB modulation. Our results suggest that during the first day post-FUS-BBB modulation, the soma of microglia increases in perimeter, which reflects more elongated or complex soma shapes.

**Figure 3.**
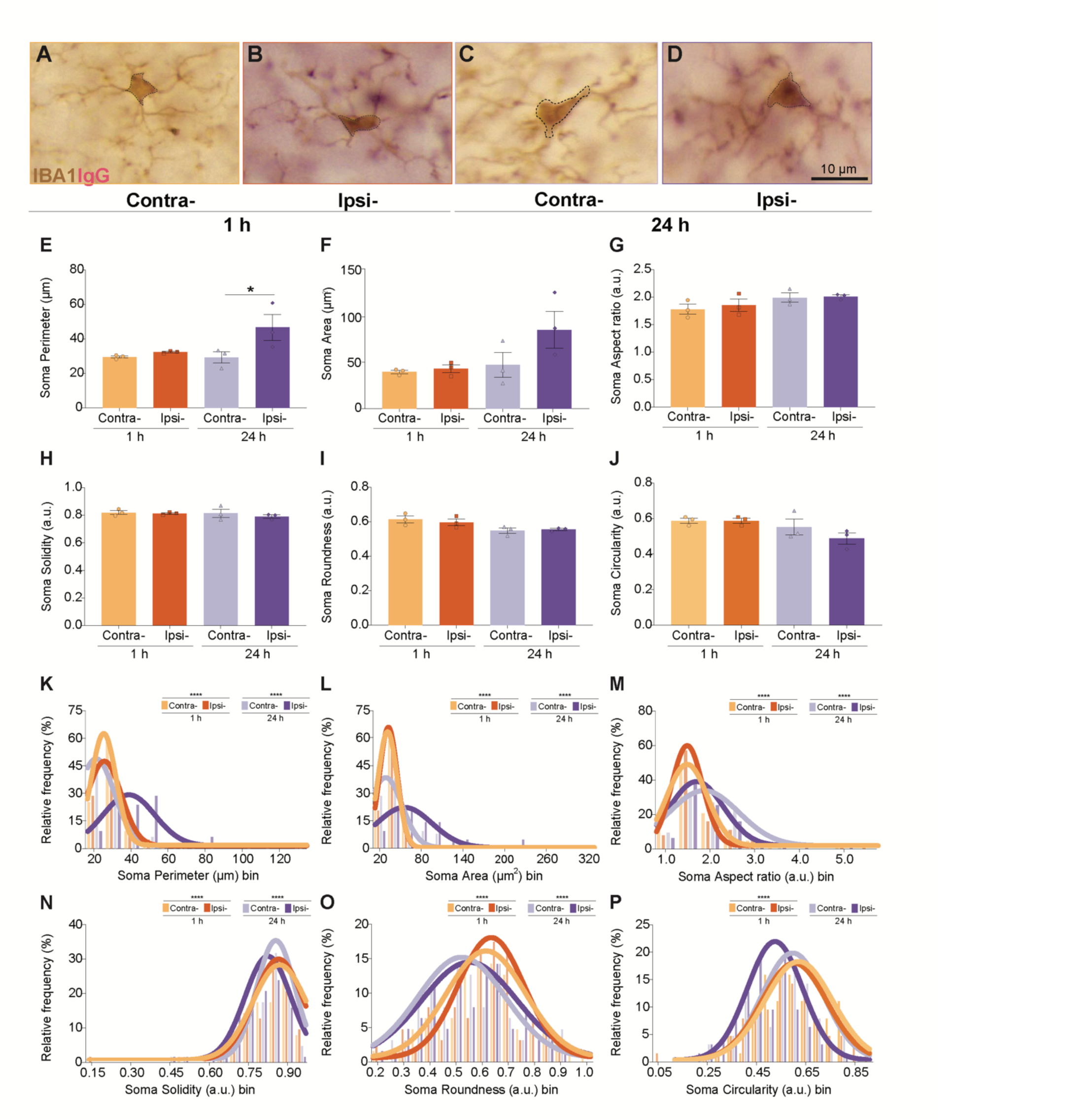
The distribution of microglial soma shape descriptors shifted one day following FUS-BBB modulation. **A-D.** Representative brightfield microscopy images taken at 40x illustrate mouse ionized calcium-binding adapter molecule (Iba)1 positive (+) cells with soma tracings (dashed line) across the contralateral (contra-) and ipsilateral (ipsi-) lacunosum moleculare (*LMol*) 1 hour (h) and 24 h following FUS-BBB. Scale bar 10 μm. **E–J.** The average soma perimeter (**E**) was significantly increased in the ipsilateral *vs* contralateral *LMol* at 24 h, according to a main effect of Hemisphere, an interaction effect of Hemisphere x Time (both not shown) and post-hoc effect for contralateral vs ipsilateral at 24 h. The mean Iba1+ soma area (**F**), aspect ratio (**G**), solidity (**H**), roundness (**I**), and circularity (**J**) did not change in the ipsilateral *LMol* at 1 h or 24 h, compared to the contralateral controls. The bar graphs show the mean, standard error of the mean, and individual data points (n = 3 animals/hemisphere/time point) analyzed by a mixed-effects 2-way analysis of variance (ANOVA) with Šídák’s multiple comparison tests. **K–P.** According to nonlinear regression models, the relative distribution of Iba1+ soma perimeter (**K**), area (**L**), aspect ratio (**M**), solidity (**N**), and roundness (**O**) was significantly distinct in the ipsilateral compared to the contralateral *LMol* at both 1 h and 24 h, though only at 1 h soma circularity (**P**). Histograms show the relative frequency and nonlinear regression analyzed via Wilcoxon comparison tests. * *p* < 0.05, ** *p* < 0.01, *** *p* < 0.001, **** *p* < 0.0001. a.u.: arbitrary unit.

Microglia present heterogeneous morphologies [4]. Average changes in morphological features represent the response of the microglial population but can potentially mask the diverse responses of individual cells. To gain insight into how FUS-BBB modulation may affect subsets of cells, we next examined the relative distribution of microglial shape descriptors using nonlinear regression models [33,34]. We found that the regressed relative distribution (Table S5) of microglial soma perimeter (Figure 3K, 1 and 24 h *p* < 0.0001), area (Figure 3L, 1 and 24 h *p* < 0.0001), aspect ratio (Figure 3M, 1 and 24 h *p*< 0.0001), solidity (Figure 3N, 1 and 24 h *p* < 0.0001) and roundness (Figure 3O, 1 and 24 h *p* < 0.0001) significantly changed in the ipsilateral *vs* contralateral *CA1 LMol* at 1 h and 24 h, though only at 1 h for microglial soma circularity (Figure 3P, 1 h *p* < 0.0001). Therefore, FUS-BBB modulation not only broadly increased microglial soma perimeter within animals, but also altered the distribution of microglial soma morphologies across the cell population.

To characterize the direction of the microglial soma morphological shifts, we analyzed the amplitude (spread) and mean (center of distribution) of each regression curve (Figure 3K-P, Table S5). Briefly, a higher curve amplitude indicates a narrower distribution of values, therefore, less variability in that morphological feature across cells. A higher curve mean reflects a shift toward larger values of the feature across the population. Correspondingly, the regressed means revealed that, in comparison to the control, microglia in the ipsilateral hemisphere deviated to larger and rounder somas with porous shapes at 1 h and 24 h following FUS-BBB (see Supplementary Methods for a detailed description). As a result, 1 h following FUS-BBB, there was a more homogenous distribution of microglial soma area and compactness, despite more variability for the distribution of soma perimeter. At 24 h, the relative distribution of all soma shape descriptors became more heterogeneous. Overall, the nonlinear regression models indicated a shift toward an increase in microglial soma size, resulting in an increasing diversity within the distribution of *LMol* soma shapes between 1 h and 24 h after BBB-FUS.

### The distribution of microglial processes shifted

In addition to affecting microglial cell bodies, the entry of blood antigens into the brain parenchyma may influence the number and structure of microglial processes, which dynamically remodel to survey various cellular compartments and the extracellular space [3]. To investigate this, we examined whether FUS-BBB alters the territory covered by microglial processes at 1 h and 24 h in the *CA1 LMol.* We traced the endpoints of Iba1+ processes to create a convex shape representing their cellular territory and, using a semi-automated pipeline, delineated masks of Iba1+ processes within each convex shape (Figure 4A-D). Similar to territory averages provided by the convex shape (in the Supplementary Table S4), the averages per animal of mask area (Figure 4E), perimeter (Figure 4F), aspect ratio (Figure 4G), circularity (Figure 4H), solidity (Figure 4I), as well as lacunarity (Figure 4J) and fractal dimension (in the supplementary Table S4), which increase with cell shape complexity [35], were not significantly different between the ipsilateral and contralateral *LMol* at 1 h and 24 h after FUS-BBB (Table S4). These findings indicate that microglial process organization overall remains unchanged at 1 h and 24 h after FUS-BBB.

**Figure 4.**
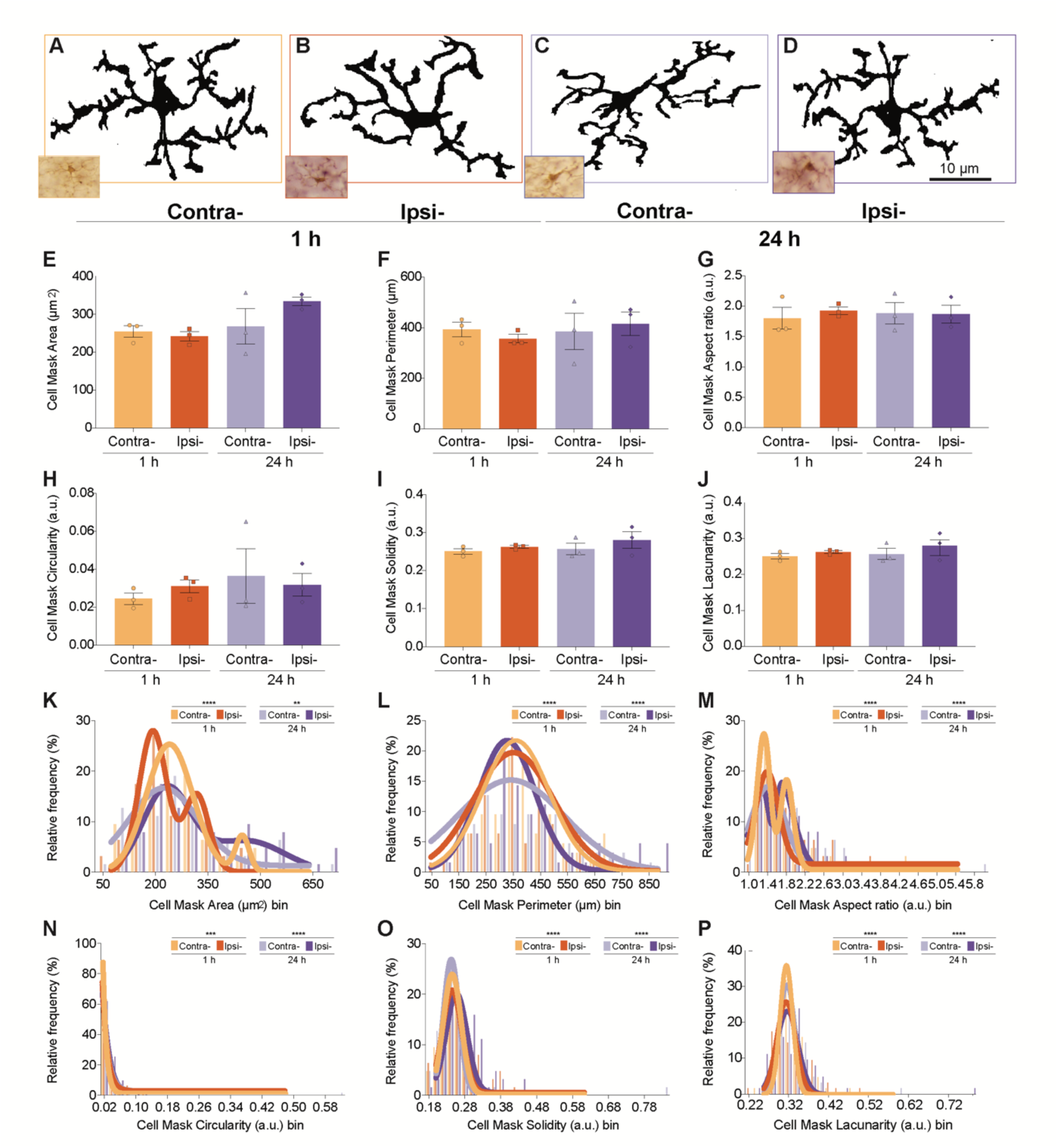
The distribution of microglial processes changed within one day of FUS-BBB modulation. **A–D.** Representative brightfield microscopy automated cell masks obtained from 40x magnification images illustrate ionized calcium-binding adapter molecule (Iba)1 positive (+) cells across the 1 h (hour) and 24 h contralateral (contra-) and ipsilateral (ipsi-) *lacunosum moleculare (LMol)* following FUS-BBB. Scale bar 10 μm. **E–J**. Iba1+ automated cell mask area (**E**), perimeter (**F**), aspect ratio (**G**), circularity (**H)**, solidity (**I**), and lacunarity (**J**) averages did not change in the ipsilateral *LMol* at 1 h and 24 h, compared to the contralateral controls. Bar graphs show the mean, standard error of the mean and individual data points (n = 3 animals/hemisphere/time point) analyzed by a mixed-effects 2-way analysis of variance (ANOVA). **K–P**. However, according to nonlinear regression models, the relative distributions of automated cell mask area (**K**), perimeter (**L**), aspect ratio **(M)**, circularity (**N**), solidity (**O)** and lacunarity **(P)** were significantly distinct in the ipsilateral compared to the contralateral control at both 1 h and 24 h. Histograms show the relative frequency and nonlinear regression analyzed via Wilcoxon comparison tests. *\* p* < 0.05, \**\* p* < 0.01, \*\**\* p* < 0.001, **** *p* < 0.0001. a.u.: arbitrary unit.

Yet, the relative distribution of microglial process shape descriptors changed among individual cells of the microglial population in response to FUS-BBB (Table S5). The nonlinear regression fits of the mask area (Figure 4K, 1 h *p =* 0.0001and 24 h *p =* 0.00), perimeter (Figure 4L, 1 and 24 h *p <* 0.0001), aspect ratio (Figure 4M, 1 and 24 h *p <* 0.0001), circularity (Figure 4N, 1 h *p =* 0.0003 and 24 h *p <*0.0001), solidity (Figure 4O, 1 and 24 h *p <* 0.0001), lacunarity (Figure 4P, 1 and 24 h *p <* 0.0001) and fractal dimension (Table S5, 1 h *p =* 0.02 and 24 h *p =* 0.00) significantly differed between the ipsilateral *vs* contralateral *LMol* at 1 h and 24 h (Table S5). To clarify the direction of these shifts within the microglial population, we referred to the spread (amplitude) and center (mean) of the distributions (Figure 4L-P, Table S5). At 1h, the microglial population appeared to adopt smaller and more regular process shapes.

At 24 h, the microglial process masks shifted back to larger, more compact and irregular shapes (see Supplementary Methods for a detailed description). Correspondingly, there was a more heterogeneous relative distribution of all microglial process descriptors at 1 h, except for the area in the ipsilateral *vs* contralateral *LMol*. The opposite was observed at 24 h, whereby the relative distribution of size and shape complexity of microglial process masks became more homogenous. Therefore, FUS-BBB resulted in a more diverse set of de-ramified (from ramified to rounder) process morphologies at 1 h and of porous morphologies at 24 h (Table S5).

We explored possible changes in microglial cell ramification further by deriving skeletonized shapes from the Iba1+ masks (Figure 5A-D), underscoring the number and length of microglial processes following FUS-BBB. The average number of branches (Figure 5E) and junctions (Figure 5F), along with the maximum branch length (Figure 5G) and the longest shortest path (Figure 5H), which all increase in value with process ramification, did not change in the ipsilateral compared to the contralateral *LMol* at 1 h and 24 h (Table S4). These findings suggest that FUS-BBB modulation does not affect overall microglial arborizations across the examined time points. However, similar to what is described above, there were noticeable shifts within the microglial population in the relative distribution of skeleton descriptors, as the nonlinear regression curves of the relative distribution of branches (Figure 5I, 1 and 24 h *p* = 0.0001), junctions (Figure 5J, 1 and 24 h *p* = 0.0001), maximum branch length (Figure 5K, 1 and 24 h *p* = 0.0001) and longest shortest path (Figure 5L, 1 h p = 0.0001, 24 h *p* = 0.0451) significantly differed between the ipsilateral vs contralateral *LMol* at 1h and 24 h (Table S5). The amplitudes and means (see Supplementary Methods for a detailed description) of the nonlinear regression models suggested that the relative distribution of microglial cells with numerous but compact branches increased at 1 h (Figure 5I-L, Table S5). By contrast, less ramified, elongated microglial cells increased at 24 h. In addition, at 1 h and 24 h, there was an overall more homogenous distribution of process properties across microglial cells within the population.

**Figure 5.**
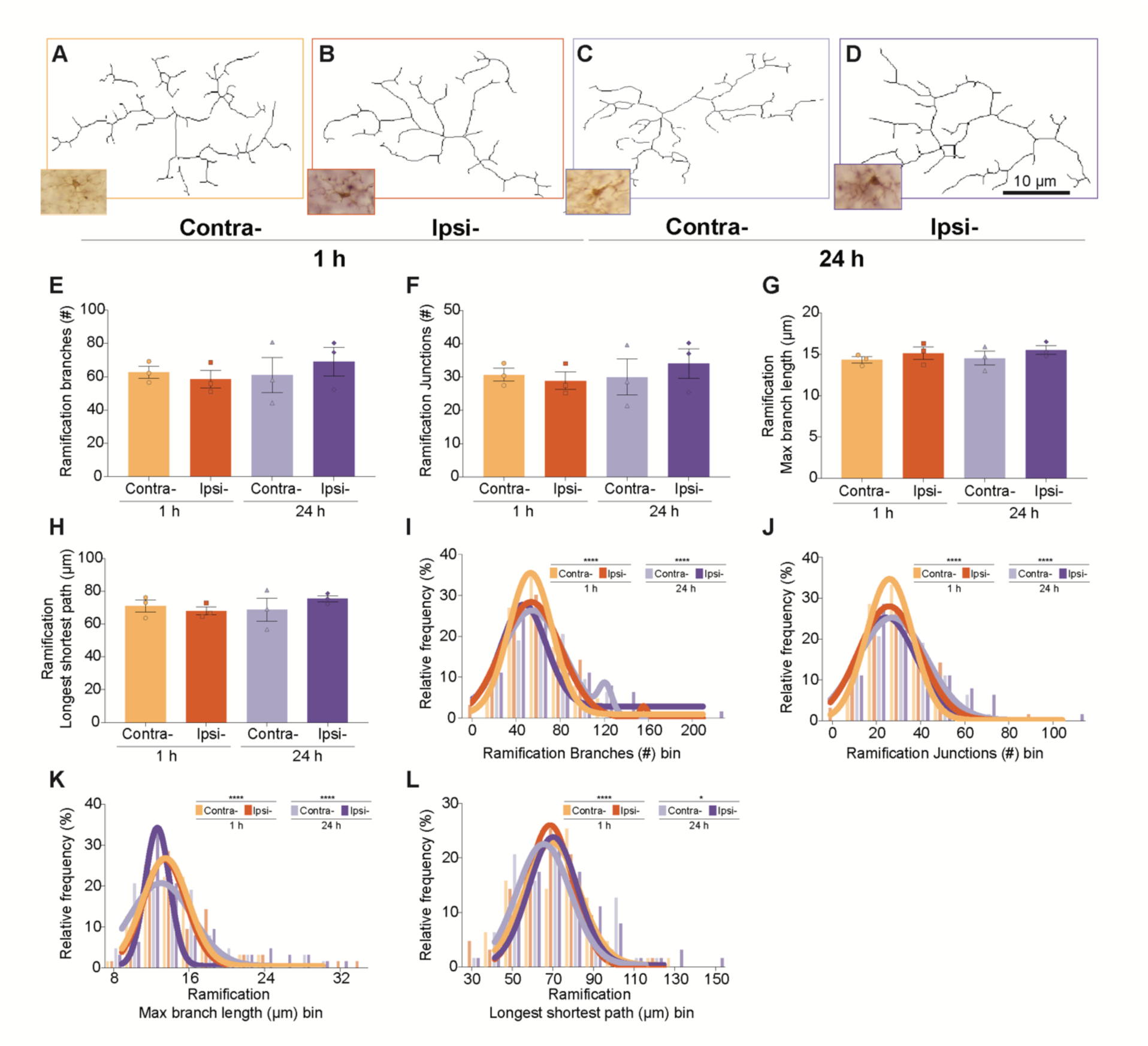
The length and number of microglial processes fluctuated 1 h and 24 h following FUS-BBB modulation. **A–D.** Representative brightfield microscopy arborizations obtained from 40x magnification images illustrate ionized calcium-binding adapter molecule (Iba)1 positive (+) cells across the 1 hour (h) and 24 h contralateral (contra-) and ipsilateral (ipsi-) *lacunosum moleculare (LMol)* following FUS-BBB. Scale bar 10 μm. **E–H.** The number of branches (**E**) and junctions (**F**), along with the maximum branch length (**G**) and longest shortest path (**H**) averages did not change in the ipsilateral *LMol* at 1 h or 24 h, compared to the contralateral controls. The bar graphs show the mean, standard error of the mean, and individual data points (n = 3 animals/hemisphere/time point) analyzed by a mixed-effects 2-way analysis of variance (ANOVA). **I–L.** By contrast, according to nonlinear regression models, the relative distributions of branches (**I**) and junctions (**J**) number, as well as maximum branch length (**K**) and longest shortest path (L) were significantly distinct in the ipsilateral *LMol* compared to the contralateral control at both 1 h and 24 h. Histograms show the relative frequency and nonlinear regression analyzed via Wilcoxon comparison tests. *\* p* < 0.05, \**\* p* < 0.01, \*\**\* p* < 0.001, **** *p* < 0.0001. a.u.: arbitrary unit.

In summary, FUS-BBB modulation is not associated with large-scale changes in microglial territory or process shape at both the 1 h and 24 h time points. However, among the cell population, there were shifts in the distribution of morphological features, which may indicate that subsets of cells change their morphology after FUS-BBB modulation. The shifts suggested a tendency to adopt bigger and more elongated soma morphologies at 1 h and 24 h following FUS-BBB. In addition, cellular architecture was remodeled into more compact territories, with stubby processes at 1 h, reverting to larger territories with fewer and simpler processes at 24 h. Overall, this resulted in a more heterogeneous distribution of microglial shapes at 1 h and less diverse process arborizations at 24 h (Figure 6).

**Figure 6.**
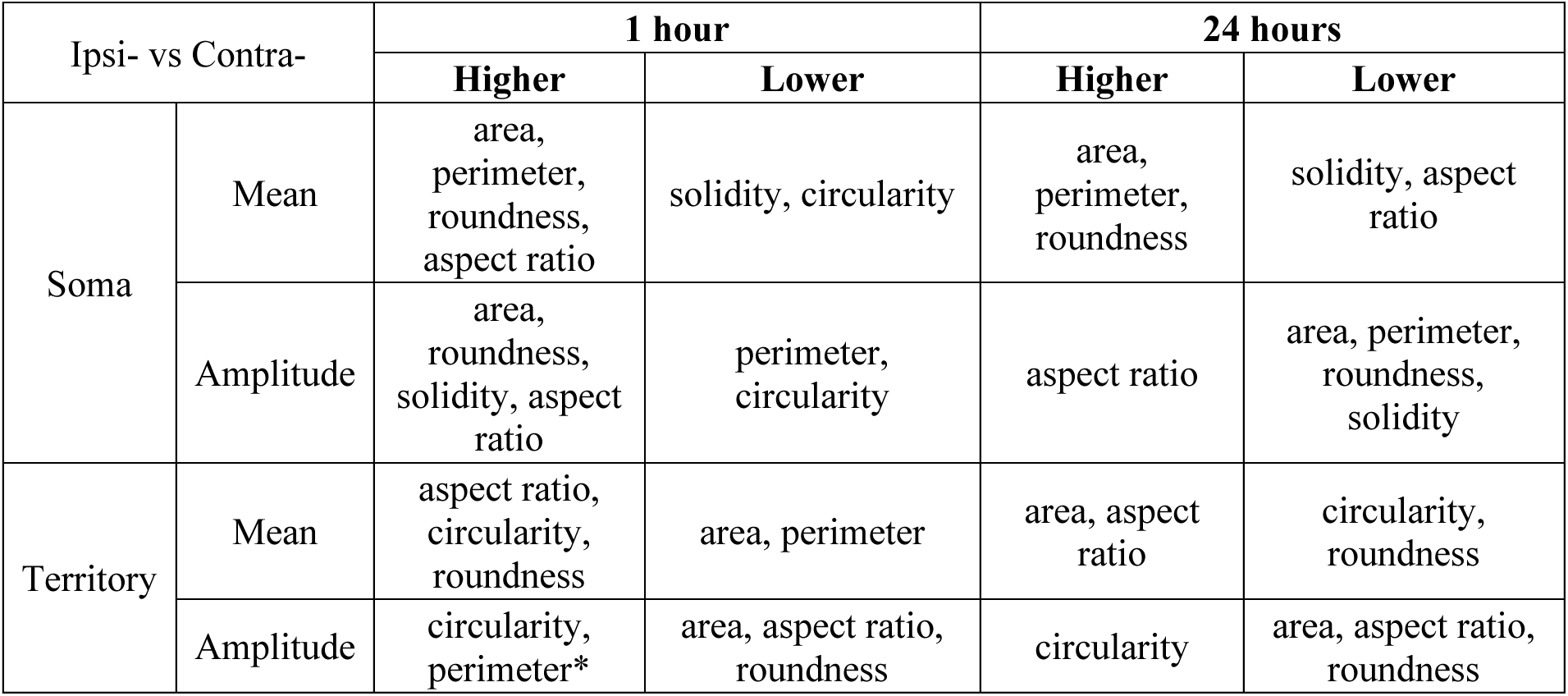
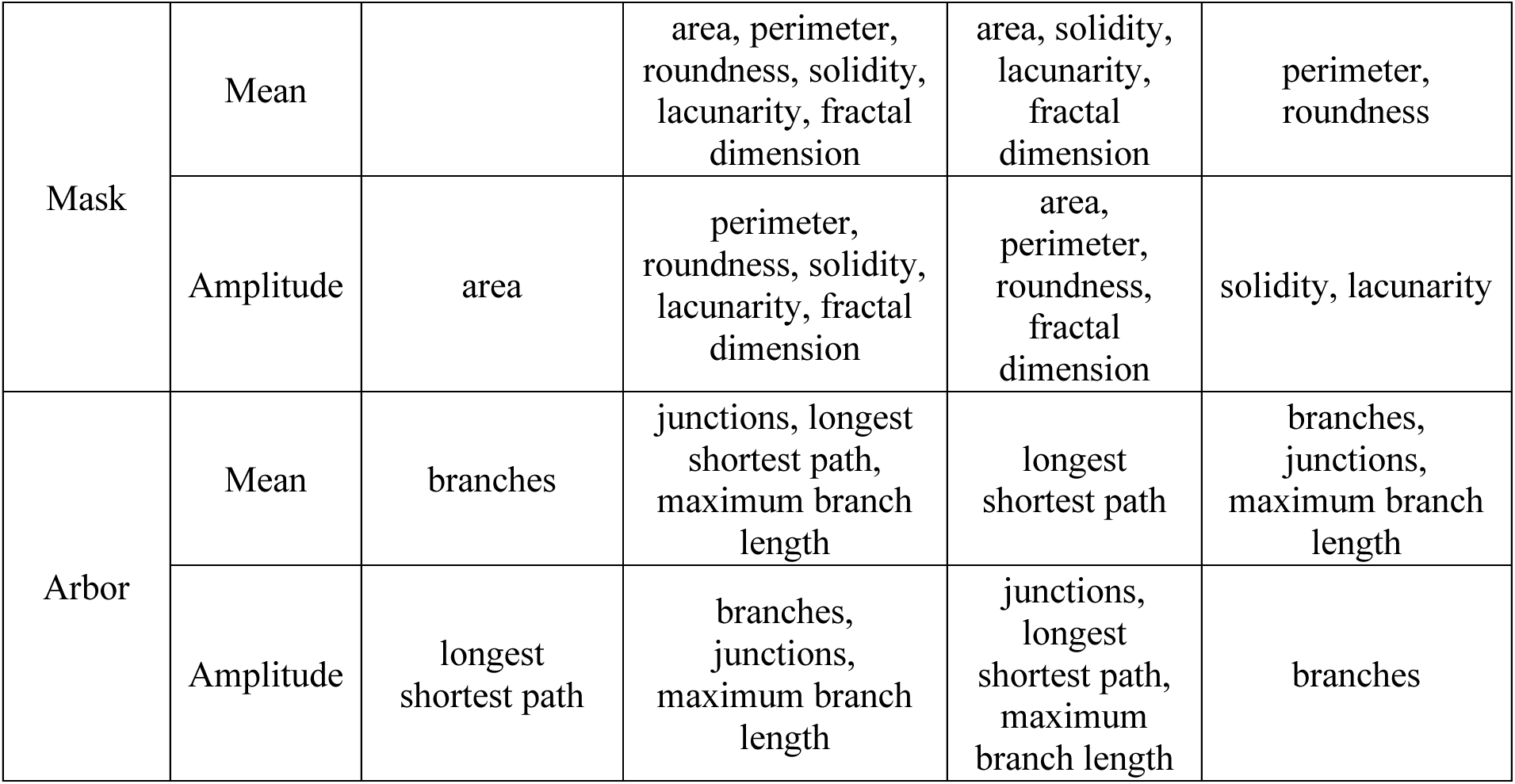
Summary of mean and amplitude nonlinear regression models in ipsilateral *versus* contralateral microglia.

Lastly, we determined whether these shifts in microglial morphology followed the extent of extravasation of blood molecules into the CNS [15]. We examined the Spearman r correlation coefficients between the optical density of IgG staining and microglial morphology parameters at 1 h and 24 h after FUS-BBB. We did not observe any significant correlations between optical density and microglial morphology (Figure S1B, Table S3), suggesting that the levels of IgG infiltration did not hold a direct relationship with shifts in microglial shape. We also examined the multidimensionality in the morphology analysis by subjecting the 30 variables of the dataset to a principal component analysis (Table S7) and evaluating which morphological parameters contributed most to the variability between ipsilateral and contralateral conditions. The analysis selected six principal components (PC), with PC1 explaining 39.23% and PC2 13.91% of the variations in the dataset (Figure S1C, Table S7). The distribution of all data points in PC1 and PC2 suggested that microglia in the ipsilateral *LMol* at 24 h contributed to most of the variability within the dataset (Figure S1C). Moreover, the loadings of variables for the PC1 and PC2 (Figure S1D) indicated that across all variables, arborization parameters such as number of branches and junctions, most strongly correlated with variation within PC1 (Table S7).

### Microglia maintained interactions with the BBB

Electron microscopy studies have provided key insights into the function of microglia, revealing their interactions with brain compartments such as synapses and the BBB [36,37]. After systemic inflammation associated with BBB leakage, several electron microscopy studies have suggested that microglia increase their proximity to blood vessels [4,38,39] and seal possible leakages [39,40]. This raises the intriguing possibility that following FUS-BBB, microglia may increase their contacts with the BBB, particularly in areas with large blood vessels, such as the *LMol* [31]. We used high-throughput chip mapping SEM to quantify the interactions between Iba1+ cells and blood vessels, including capillaries, veins and arteries (Figure 7A-D). Intriguingly, we observed a single Iba1+ process enclosed between a capillary in the ipsilateral *LMol* at 1 h (Figure 7B). This process may be probing or sealing the modulated BBB, as previously suggested [39,40]. However, we did not find significant differences in the average distance between microglia and blood vessels (Figure 7E, Table S8) at 1 h and 24 h after FUS-BBB, nor in the average number of microglia physically apposed to vessels (<150 nm) (Table S8). It is possible that, instead of broadly increasing contacts or proximity to blood vessels, microglia probed different elements of the neurovascular unit following FUS-BBB modulation [15]. We verified first whether FUS could influence microglial cell body interactions with the basement membrane (Figure 7A-D), a form of extracellular matrix that anchors cells in the BBB and supports signal transduction [15]. There was a significant interaction between Hemisphere and Time point in microglial contacts with the basement membrane (Figure 7F, interaction effect of Hemisphere x Time, F = 12.31, *p* = 0.02), but pairwise comparisons did not reach significance.

**Figure 7.**
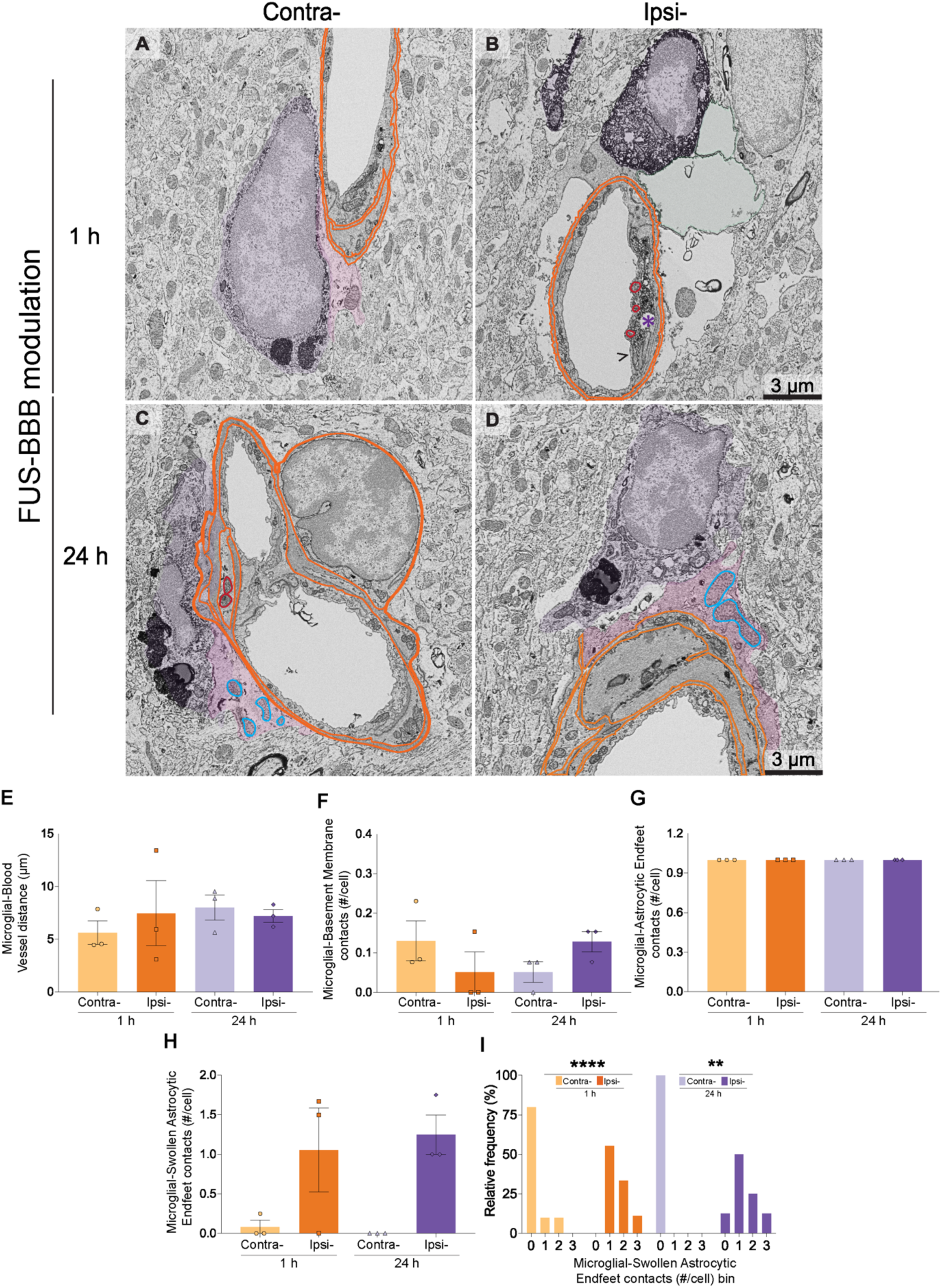
FUS-BBB modulation increased the frequency of microglial contact with swollen astrocytic endfeet. **A-D**. Representative scanning electron microscopy (SEM) images of ionized calcium-binding adapter molecule (Iba)1 positive (+) cell bodies (purple) and process (purple asterisk), alongside blood vessels in the ipsilateral (ipsi-) and contralateral (contra-) *stratum lacunosum moleculare* (*LMol*) at 1 hour (h) and 24 h after FUS-BBB. Microglial contacts with endothelial cells and astrocytic endfeet contain mitochondria, pseudocolored in red and blue respectively. Scale bar 3 μm. **E–H**. The average distance between microglia and blood vessels (**E**), the number of contacts between microglia and the basement membrane (**F,** orange), astrocytic endfeet (**G**, pink) and swollen astrocytic endfeet (**H**, green) did not differ between the ipsilateral and contralateral *LMol* of mice at 1 h and 24 h after FUS-BBB modulation, despite an interaction effect between Hemisphere and Time for basement membrane contacts and a Hemisphere effect for swollen endfeet contacts (both not shown). The bar graphs show the mean, standard error of the mean, and individual data points (n = 3 animals/hemisphere/time point) analyzed by a mixed-effects 2-way analysis of variance (ANOVA). **I**. By contrast, the relative number of contacts between microglia and swollen astrocytic endfeet significantly increased in the ipsilateral compared to the contralateral *LMol* at 1 h and 24 h after FUS-BBB modulation. Relative frequency bar graphs analyzed via Fisher’s exact tests. *\*p* < 0.05, \**\*p* < 0.01, \*\**\*p* < 0.001.

At the neurovascular unit, microglia also frequently interact with astrocytic endfeet, which provide a crucial link between neuronal and endothelial cell function to modulate blood flow [15]. We observed numerous interactions between microglial cell bodies and astrocytic endfeet (Figure 7A-D). However, these did not change in the *LMol* across contralateral and ipsilateral hemispheres at 1 h and 24 h after FUS-BBB (Figure G). Astrocytic endfeet can become swollen in health and disease, increasing in size and showing a clear cytoplasm enclosed by a plasma membrane [41]. Accordingly, we observed swollen endfeet in both ipsilateral and contralateral *LMol* at 1 h and 24 h (Figure 7B). There was a significant effect of the Hemisphere on the average number of contacts microglia made with swollen astrocytic endfeet (Figure 7H, main effect of Hemisphere f =16.48, *p* = 0.02), but pairwise comparisons at 1 h and 24 h did not reach significance. We compared instead the relative frequency of contacts between microglia and swollen endfeet and found they were significantly higher in ipsilateral *vs* contralateral *LMol* at 1 h and 24 h (Figure 7I, 1 h *p =* 0.0007, 24 h *p =* 0.0101, Table S9). These findings suggest that FUS-BBB is not associated with a general increase in microglia-BBB interactions but rather a specific recruitment towards vessels with swollen astrocytic endfeet.

We next investigated whether vessel proximity and diameter play a role in modulating microglial-BBB interactions through a Spearman r correlation, including both time points (Figure S2A, Table S12). Blood vessel distance significantly and negatively correlated with interactions between microglia and the basement membrane (Figure S2A, r = -0.4722, *p* = <0.0001), homeostatic (Figure S2A, r = -0.7330, *p* =<0.0001), swollen astrocytic endfeet (Figure S2A, r = -0.3126, *p* = 0.0019) and astrocytic endfeet containing mitochondria (Figure 7C-D, Figure S2A, r = -0.6213, *p* =0.0000), proposed to attract microglia [40]. Moreover, we observed a significant positive correlation between vessel area and interactions between microglia and astrocytic endfeet (Figure S2A, r = 0.3624, *p* = <0.0001), swollen astrocytic endfeet (Figure S2A, r = 0.3072, *p* = 0.0001), the basement membrane (Figure S2A, r = 0.1793, *p* = 0.0276), as well as endothelial cells (Figure 7A-B, Figure S2A, r = 0.1800, *p* = 0.0270) and astrocytic endfeet (Figure S2A, r = 0.3151, *p* = 0.0001) containing mitochondria. These results suggest that, following FUS-BBB modulation, microglial cell bodies that are distant from vessels are less likely to interact, while larger vessels are more likely to receive microglial contacts.

### Microglia reduced contacts with pre-synaptic elements

Microglia modulate synaptic plasticity through many mechanisms, including partial engulfment of pre-synapses, known as trogocytosis [42]. FUS-BBB modulation has also been proposed to trigger synaptic plasticity; however, the mechanisms remain to be established [15]. Using chip mapping SEM, we verified whether FUS-BBB modulation alters the ultrastructural interactions between microglia and pre-(Figure 8A,B) and post-synaptic (Figure 8A,B,D) elements, as well as extracellular space pockets (Figure 8A,C), required for microglial motility and neuronal remodeling [43,44]. We identified that time modulated the average number of pre-synaptic elements contacted by microglial cell bodies (Figure 8E, main effect of Time, F = 33.0600, *p =* 0.00), but pairwise comparisons did not reach significance, suggesting the contacts between pre-synaptic elements and microglia changed over the 24-hour period following FUS-BBB modulation. When we looked at the cell population, there was a significant difference in the relative distribution curves of microglia-pre-synaptic contacts between contralateral and ipsilateral *LMol* at 1 h and 24 h following FUS-BBB (Figure 8F 1 and 24 h *p =* <0.0001). Accordingly, the nonlinear regression models suggested that the relative distribution of microglial contacts with pre-synaptic elements reduced at both time points, resulting in a more homogeneous distribution of contacts at 1 h and the reverse effect at 24 h (see Supplementary Methods for a detailed description, Table S10). We did not find significant differences in the average number of contacts between microglia and other neuronal compartments, such as post-synaptic elements, neuronal cell body (satellite cells), myelinated axons and degenerating myelin (Table S8). Despite no significant difference in the average number of contacts between microglial cell bodies and extracellular space pockets (Figure 8), there was a reduced frequency of contacts in the ipsilateral *vs* contralateral *LMol* at 24 h after FUS-BBB (Figure 8H, *p =*0.0040, Table S9), with no differences for pockets containing debris or presenting digestion (Table S8). Overall, our findings suggest that the microglial population shifts to reduce the frequency of contacts between microglia, pre-synaptic elements and extracellular space pockets, which might reduce the rates of trogocytosis and modulate synaptic plasticity after FUS-BBB modulation [44].

**Figure 8.**
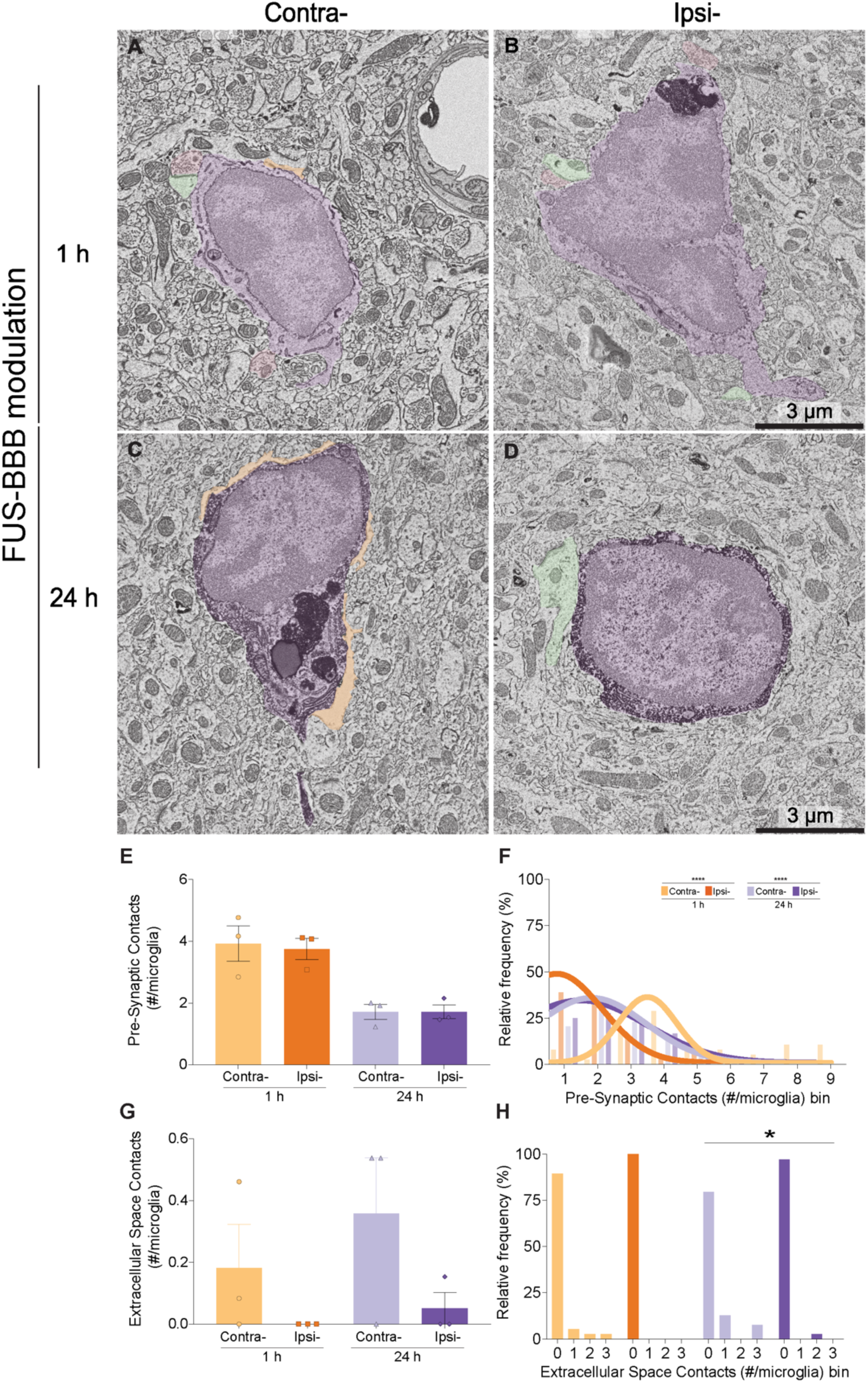
**Microglia interacted less frequently with pre-synaptic elements and extracellular space pockets following FUS-BBB modulation. A–D**. Representative scanning electron microscopy (SEM) images of ionized calcium-binding adapter molecule (Iba)1 positive (+) cells (purple) and blood vessels in the ipsilateral (ipsi-) and contralateral (contra-) *stratum lacunosum moleculare* (*LMol*) at 1 hour (h) and 24 h after FUS-BBB. Microglial contacts with pre-synaptic element (**A, B,** pink), post-synaptic element (**A**, **B, D,** green) and extracellular space pockets (**C**, beige). Scale bar 3 μm. **E**. The average number of contacts between microglia and pre-synaptic elements was significantly modulated by Time (main effect not shown), suggesting a reduction in contacts during the 24 h after FUS-BBB modulation. The bar graph shows the mean, standard error of the mean, and individual data points (n = 3 animals/hemisphere/time point) analyzed by a mixed-effects 2-way analysis of variance (ANOVA) with Šídák’s multiple comparisons tests**. F**. According to nonlinear regression modeling, the relative distribution of ultrastructural interactions between microglia and pre-synaptic elements significantly differed between the ipsilateral and contralateral *LMol* at 1 h and 24 h after FUS-BBB modulation. Histograms show the relative frequency, and nonlinear regression analyzed via Wilcoxon comparison tests. **G**. The average number of ultrastructural interactions with extracellular space pockets did not differ for microglial cell bodies examined at 1 h or 24 h after FUS-BBB modulation in the ipsilateral *vs* contralateral *LMol*. **H**. However, at 24 h, the relative frequency of contacts with extracellular space per microglial cell body decreased in the ipsilateral *LMol* compared to the contralateral *LMol* at 24 h. Histograms show the relative frequency analyzed by Fisher’s exact tests. *p < 0.05, **p < 0.01, ***p < 0.001. #: number.

### Markers of microglial oxidative stress increased in a subset of cells

The ultrastructural state of microglial organelles, such as endoplasmic reticulum (ER)/Golgi and mitochondria (Figure 9A), can indicate elevated oxidative stress following BBB permeability [35]. In microglial cell bodies, we examined ultrastructural alterations in organelles typically associated with metabolic demand, including ER/Golgi dilation and mitochondria elongation (Figure 9B), ER/Golgi and mitochondrial density (Figure 9C), as well as damage, i.e., dystrophic mitochondria (Figure 9D) [35] following FUS-BBB using chip mapping SEM in the ipsilateral *vs* contralateral *CA1 LMol*. There were no significant differences in microglial density of ER/Golgi (Figure 9E), homeostatic (Figure 9F), elongated (Figure 9G) and dystrophic mitochondria (Figure 9H) at 1 h and 24 h, despite an opposing modulatory effect of Hemisphere on the number of homeostatic (Figure 9F, main effect of the Hemisphere F = 17.3200, *p =* 0.0141) and dystrophic (Figure 9H, main effects of the Hemisphere F = 5.5910, *p =* 0.0456) mitochondria (Table S8). Moreover, across the microglial cell population, the nonlinear regression curves of ipsilateral *vs* contralateral distributions of ER/Golgi (Figure 9I, 1 and 24 h *p =* <0.0001), homeostatic mitochondria (Figure 9J, 1 and 24 h *p =* <0.0001) and elongated mitochondria (Figure 9K, 1 and 24 h *p =* <0.0001) were significantly different at 1 h and 24 h (Table S10).

**Figure 9.**
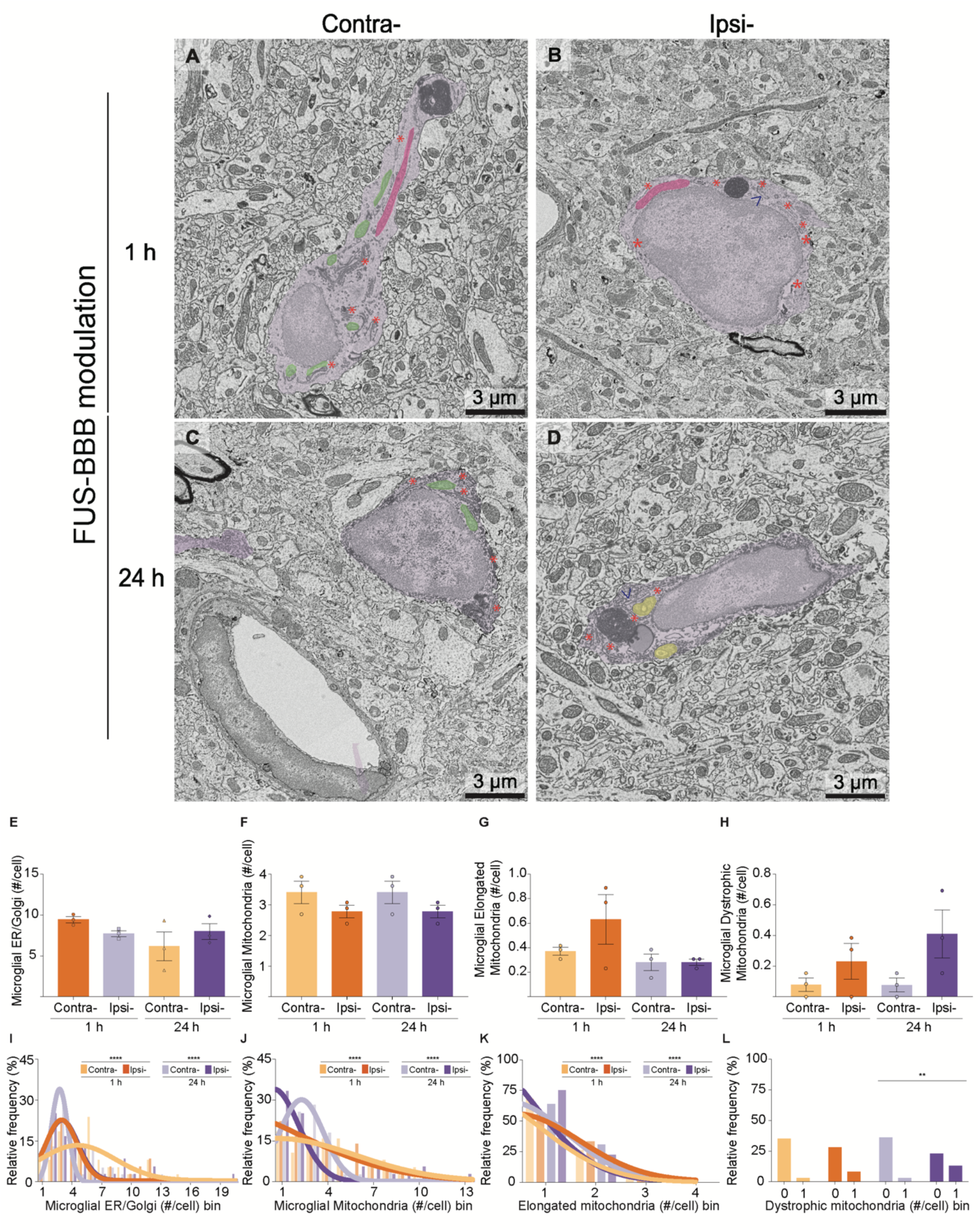
Microglia altered their mitochondria and endoplasmic reticulum following FUS-BBB modulation. **A–D.** Representative scanning electron microscopy (SEM) images of ionized calcium-binding adapter molecule (Iba)1 positive (+) cells (purple) and blood vessels in the ipsilateral (ipsi-) and contralateral (contra-) *stratum lacunosum moleculare* (*LMol*) at 1 hour (h) and 24 h after FUS-BBB. Pictures highlight microglial homeostatic mitochondria (green) and endoplasmic reticulum/golgi apparatus (ER/Golgi, orange asterisk) cisternae (**A, C**), elongated mitochondria (**A, B**, pink), dilated ER/Golgi (**B, D**, blue arrowhead), and dystrophic mitochondria (**D**, orange). Scale bars: 3 μm. **E–-H**. The average number of microglial ER/Golgi cisternae (**E**), mitochondria (**F**), elongated mitochondria (**G**) and dystrophic mitochondria (**H**) did not change at 1 h and 24 h after FUS-BBB modulation in the ipsilateral and contralateral *LMol,* despite a main effect of Hemisphere for homeostatic and dystrophic mitochondria (both not shown). The bar graphs show the mean, standard error of the mean, and individual data points (n = 3 animals/hemisphere/timepoint) analyzed by a mixed-effects 2-way analysis of variance (ANOVA). **I-K**. According to the nonlinear regression models, the relative frequency of ER/Golgi (**I**), mitochondria (**J**) and elongated mitochondria (**K**) differed between varied in the ipsilateral *vs* contralateral *LMol* at 1 h and 24 h. **L**. Histograms show the relative frequency and nonlinear regression analyzed via Wilcoxon comparison tests. The number of dystrophic mitochondria per cell was significantly distinct between ipsilateral and contralateral *LMol* at 24 h. Histograms show the relative frequency analyzed via Fisher’s exact tests. *\*p* < 0.05, \**\*p* < 0.01, \*\**\*p* < 0.001. #: number.

According to the amplitude and means of the regression models, at 1 h but not 24 h in the ipsilateral *LMol,* the relative distribution of microglial cell bodies with higher metabolic demands increased within the population (see Supplementary Methods for a detailed description). Such shifts resulted in an increasingly heterogeneous ER/Golgi distribution across microglial cell bodies at 1 h and 24 h, with the reverse effect for homeostatic and elongated mitochondria. We also observed a higher frequency of microglial cell bodies with dystrophic mitochondria at 24 h in the ipsilateral *LMol versus* contralateral *LMol* after FUS-BBB (Figure 9L, *p =* 0.0040, Table S9). Overall, our findings suggest that FUS-BBB modulation is associated with increased microglial oxidative stress, resulting in a higher number of cells with increased ER/Golgi activity and mitochondrial alterations in a subset of cells.

### Lysosomal activity decreased in microglia

FUS-BBB modulation has been associated with increased phagolysosomal activity, based on increased microglial inclusion of pathological proteins [17]. To provide further insights, we used chip mapping SEM to quantify the number of empty and filled phagosomes, autophagosomes and primary (Figure 10A), secondary (Figure 10B), and tertiary (Figure 10C), lysosomes within microglial cell bodies (Figure 10D) in the contralateral and ipsilateral *LMol* at 1 h and 24 h following FUS-BBB. We observed significantly fewer primary lysosomes per microglial cell body at 24 h (Figure 10E, interaction effect of Hemisphere x Time F = 12.7200 and *p =* 0.0235, post-hoc contralateral vs ipsilateral at 24 h *p =* 0.0368 and t = 3.8340). There was a significant interaction between Time and Hemisphere for secondary lysosomes (Figure 10F, interaction effect of Hemisphere x Time F = 8.6430 and *p =* 0.0424), despite no significant difference in pairwise comparisons, suggesting our analysis may have been underpowered in this dataset. We also did not observe any significant variation in tertiary lysosomes (Figure 10G) between the ipsilateral and contralateral *LMol* across the time points. However, there were fewer microglial cell bodies with primary (Figure 10H, *p =* 0.0259) and secondary lysosomes at 24 h (Figure 10I, *p =* 0.0379), but no impact on tertiary lysosomes (Figure 10J, Table S9). These findings suggest that primary and secondary microglial lysosomal activity decreased at 24 h after FUS-BBB. This was a specific effect on lysosomal digestion and not phagocytosis, as we did not find significant differences for the average number and relative frequency of empty and filled phagosomes and autophagosomes (Table S8-9).

**Figure 10.**
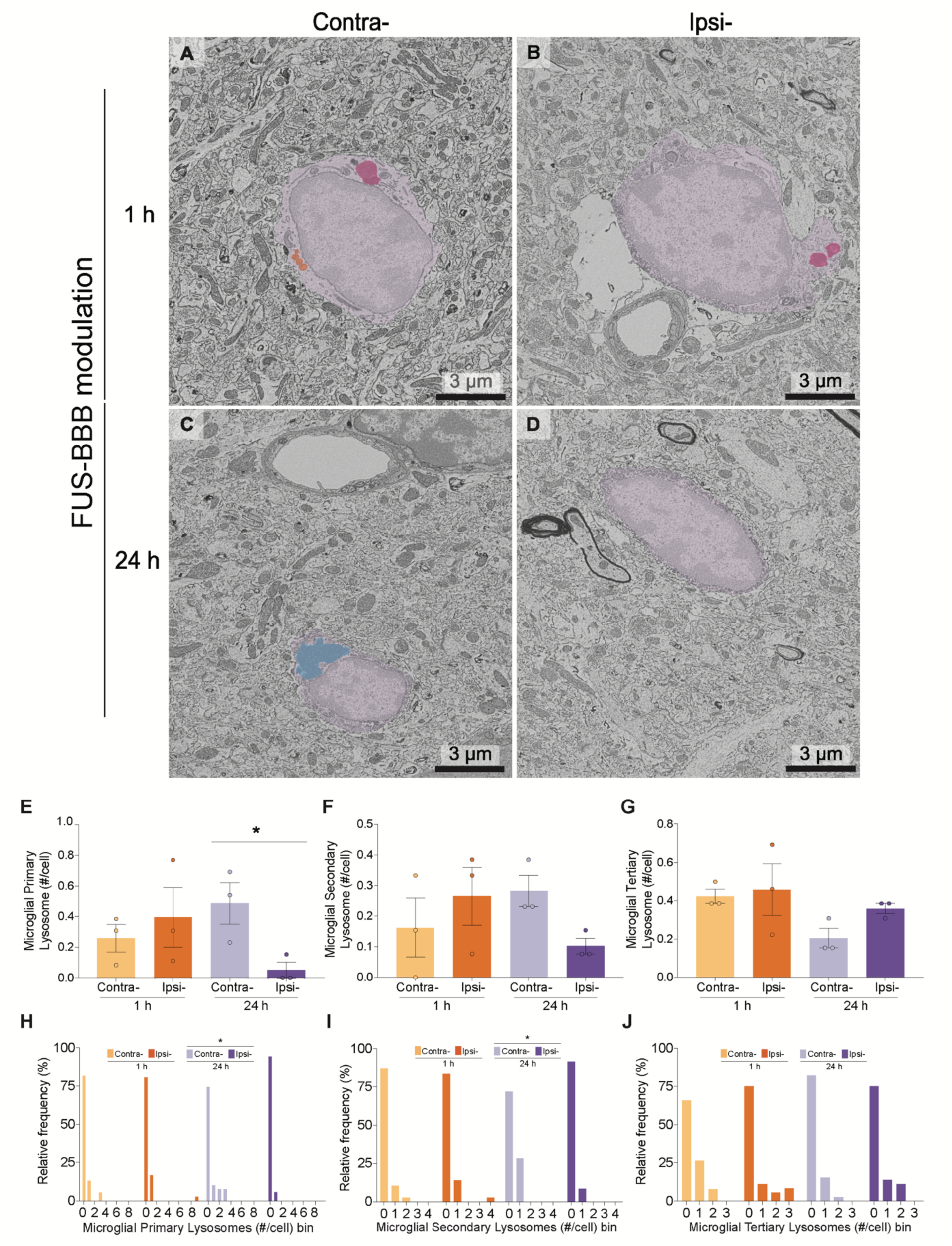
Microglia had fewer primary lysosomes following FUS-BBB modulation. **A–D.** Representative scanning electron microscopy (SEM) images of ionized calcium-binding adapter molecule (Iba)1 positive (+) cells (purple) and blood vessels in the ipsilateral (ipsi-) and contralateral (contra-) *stratum lacunosum moleculare* (*LMol*) at 1 hour (h) and 24 h after FUS-BBB. Pictures highlight primary (**A**, orange), secondary (**B**, pink) and tertiary (**C**, blue) lysosomes and microglial cell bodies without visible lysosomes (**D)**. Scale bars: 3 μm. **E**. The average number of primary lysosomes was significantly decreased in microglial cell bodies of the ipsilateral *vs* contralateral *LMol* at 24 h after FUS-BBB, according to both an interaction effect between Time and Hemisphere (not shown) and post-hoc comparison. **F**. A significant interaction effect (not shown) that did not pass pairwise comparisons was observed for the average of secondary lysosomes between ipsilateral and contralateral *LMol*. **G**. The average number of tertiary lysosomes did not change between ipsilateral and contralateral *LMol* at 1 h and 24 h after FUS-BBB. The bar graphs show the mean, standard error of the mean, and individual data points (n = 3 animals/hemisphere/time point) analyzed by a mixed-effects 2-way analysis of variance (ANOVA) with Šídák’s multiple comparisons tests. **H–J**. The relative frequency of primary (**H**) and secondary lysosomes (**I**) were also significantly different between ipsilateral and contralateral *LMol* at 24 h after FUS-BBB, with no changes in tertiary lysosomes (J). Histograms show relative frequencies analyzed via Fisher’s exact tests. *\*p* < 0.05, \**\*p* < 0.01, \*\**\*p* < 0.001.

Modifications in microglial contacts with blood vessels and the neuropil can occur simultaneously to changes in their organelles. To address multidimensionality within the ultrastructure dataset, we explored how each of 30 ultrastructural variables related to one another and with FUS-BBB using PCA (Table S11). The analysis selected three PC, with PC1 explaining 13.78% and PC2 11.05% of the variations in the dataset, indicating around 25% of the total variability in the dataset can be explained by the effects of combined ultrastructural variables (Table S11). The distribution data points in PC1 and PC2 were more dispersed in the ipsilateral *LMol* at 24 h, compared to other groups, suggesting that most variability in the ultrastructure dataset came from microglial changes at 24 h in the targeted area (Figure S2C). Moreover, the loadings of variables for PC1 and PC2 indicated that across all variables, parameters such as number of contacts between microglial cell bodies and astrocytic endfeet, swollen astrocytic endfeet, the basement membrane, along with alterations in ER/Golgi and mitochondria, most strongly correlated with variation within PC1 and PC2 (Figure S2D and Table S11).

## Discussion

Several reports have looked at baseline microglial responses starting a day after FUS-BBB modulation [26,45,46,46,47], but similar evidence from the earlier hours of FUS-induced BBB permeability is still limited [15,25,47]. Our study explored whether microglial properties change early on following FUS-BBB. We leveraged hippocampal FUS-BBB to investigate microglial density, distribution, morphology, and ultrastructure at peak BBB permeability and after its resolution at 1 h and 24 h, respectively. We found that microglia generally do not change their density and distribution in the early stages after FUS-BBB. Cellular morphology was largely stable, with subsets of cells shifting towards shapes that were more compact and stubbier at 1 h, as opposed to more extensive and simpler at 24 h. Microglia likely increased surveillance of blood vessels presenting astrocytic swelling at both time points, while microglial contacts with pre-synaptic elements and extracellular space pockets tended to decrease at 24 h. In parallel, microglial cell bodies exhibited increased dystrophic mitochondria and reduced primary lysosomes at 24 h, suggesting an increased metabolic demand and disrupted digestive activity. Our findings indicate that FUS-induced BBB permeability does not drive early global changes in microglial surveillance and phagocytosis, but induces subsets of microglia to shift their shape, cell-cell interactions and cellular activity.

To our knowledge, this is the first study to characterize microglial morphology early upon FUS-BBB [15]. Only the microglial soma perimeter was significantly increased in the treated hemisphere at 24 h after FUS-BBB. This finding contrasted with the elevated microglial soma area previously observed in cells adjacent to the targeted hippocampi a week after FUS-BBB [18]. An early increase in perimeter not accompanied by changes in area could indicate a more elongated soma shape reminiscent of rod microglia [48], which are proposed to shield dendrites and axons, notably after traumatic brain injury in rats [49]. It could also be an indication of membrane ruffling associated with cellular motility, receptor internalization and other activities [50]. Indeed, there was an increase in the relative distribution of larger, rounder and more porous microglial somas at 1 h and 24 h, based on nonlinear regression. Combined with the increased heterogeneity in the relative distribution of most microglial soma shape descriptors within the population at 24 h, we speculate that individual microglia could diversify their cell body shape, some ultimately adopting a more elongated or complex morphology 24 h after FUS-BBB modulation.

Microglia can also use their processes to shield damage; only 30 min after a BBB lesion, cortical vessel-associated microglia were seen extending their processes to reduce BBB permeability in mice [24]. Our findings suggest that FUS-BBB does not drive immediate and global changes in microglial ramification.

However, microglia did shift their stubby processes at 1 h to larger, fewer and simpler processes at 24 h. Selective modulation of arborization aligns with early observations that microglia constantly modify their morphology, a process which can be intensified by the distance from and severity of BBB damage [51]. The shift towards de-ramification that we observed could support the rod-like state mentioned before; however, it is also most likely that these cells adopt more than one type of arbor morphology in response to FUS-BBB [3]. Cells transitioning from a ramified to an ameboid microglial morphology after lesion generally do so step-wise, with the withdrawal of processes, followed by soma enlargement and process protrusion [52]. Hence, the heterogeneous relative distribution of process shape descriptors we observed within the microglial population at 1 h may also reflect a push to maintain the diverse, specialized roles of microglia, in parallel to a partial transition toward ameboid cells, which may serve a protective role in the parenchyma [15,52].

Following previous evidence of microglia dynamically approaching blood vessels [4,38,39], and potentially sealing BBB leakages [39,40], we did not identify significant differences in the average distance or number of microglial cell bodies associated with blood vessels at 1 h and 24 h after FUS-BBB. Similarly, our light microscopy findings suggested the distribution of microglia did not change, despite a main effect of time in modulating the overall distance between cells, or NND. A significantly reduced NND of hippocampal microglia was previously observed in mice one week after FUS-BBB modulation [14]. Moreover, single-cell sequencing analysis of mouse hippocampal microglia at 24 h after FUS-BBB modulation revealed a molecular profile indicative of cell proliferation [20]. It is plausible that changes in microglial density and distribution after FUS-BBB take longer than 24 h. Contrary to rapid process movements [24], rodent microglial cell bodies are much less mobile, both at baseline and after BBB lesion [51], with only 7.5% of microglial cell bodies shifting >5 μm toward vessels over several days [53]. Thus, we argue that while microglial cell bodies do not migrate toward vessels in the 24 h after FUS-BBB, they may do so later, as identified in non-human primates at a 48 h time point [46].

In addition, it is possible that changes in microglial distribution occur in some proportion to plasma extravasation into the CNS [30]. In rodent studies where microglial cell bodies quickly approached vessels, e.g. after daily injection of lipopolysaccharide [39] or stroke [40], the BBB suffered more intense insults compared to a single FUS-BBB [6]. We identified a correlation between IgG optical density and NND 1h and 24 h after FUS-BBB, suggesting elevated BBB permeability push microglia closer to one another or to damaged areas. Plasma extravasation, in turn, may differ based on vessel type, as arterioles have been proposed to be more active vesicular transport by FUS-BBB than capillaries and venules [8]. Indeed, of all hippocampal *CA1* strata, the *LMol*, with its abundant large vessels [54], was the main region to visually show endogenous IgG at 1 h and 24 h. We observed a positive correlation between vessel area and interactions between microglia and BBB elements. Moreover, we observed a significantly higher relative frequency of microglial contacts with swollen astrocytic endfeet in the ipsilateral *vs* contralateral *LMol* at 1 h and 24 h. Previously, FUS targeted at brain tumours was shown to increase astrocytic endfeet swelling in rats [55]. However, swollen endfeet also occur naturally in the *LMol,* that is, in healthy, non-treated mice [43]. Our findings suggest that larger vessels bearing swollen endfeet are likely to recruit microglia after FUS-BBB, warranting further 3-dimensional electron microscopy analyses, which can verify progressive microglial cell body and process movement toward vessels and clarify contacts between microglia and basement membrane, which were not clear in our 2-dimensional dataset.

Increased recruitment of microglia towards vessels could hinder physiological surveillance of synapses and help explain the reduction in contacts between microglial cell bodies and pre-synaptic elements we observed for subsets of cells in the targeted *LMol*. Indeed, shifts in microglial morphology towards rod or ameboid cells are typically associated with reduced surveillance [36]. Functionally, decreased microglia-pre-synaptic interactions may prevent trogocytosis, a mechanism by which microglia partially engulf pre-synapses and extracellular matrix components to stimulate dendritic spine filopodia formation [42,56].

Accordingly, we observed decreased the frequency of microglial contacts with extracellular space pockets in the ipsilateral compared to contralateral *LMol* at 24 h after FUS-BBB, raising the possibility that FUS-BBB modulation has immediate effects on synaptic plasticity by reducing trogocytosis.

Following stereotaxic injection of plasma into the brain, microglia increased their expression of genes related to oxidative stress [30]. Penetration of plasma molecules into the CNS following FUS-BBB modulation may also impact the metabolism in microglia, albeit at different degrees, due to the restrictions of plasma components that could enter the brain following FUS/and with lack of vessel damage as in stereotaxic injections [15]. We found that microglial mitochondrial damage was more frequent in the ipsilateral *vs* contralateral *LMol* 24 h after FUS-BBB, that is, while the number of microglial homeostatic mitochondria decreased with time in the ipsilateral hippocampi, the number of dystrophic microglia increased, as did their frequency per cell. This was supported by nonlinear regression models, where the distribution of microglial cells with ultrastructural markers of higher metabolic demands increased in the ipsilateral *LMol* at both time points. Accordingly, elevated [18F] DPA-714 binding to the translocator protein, non-selectively present in microglial mitochondria and typically upregulated with inflammation, was identified at 24 h after FUS-BBB in rat hippocampi [45]. In our study, indications of mitochondrial stress were accompanied by significantly fewer immature lysosomes, but stable phagosome density at 24 h in the ipsilateral *LMol*. These findings suggest that microglia maintain regular phagocytic intake but exhaust cellular metabolism and digestive activity early after FUS-BBB modulation. Sequencing of mouse hippocampal microglial 1 and 3 days following FUS-BBB modulation indicated an enrichment for lysosomal and phagocytic pathways [20]. Despite not observing changes in microglial phagosomal content, reduced digestion early after FUS-BBB modulation could cause a build-up of parenchymal debris, stimulating phagocytic activity days after, as observed through colocalization of microglial markers and misfolded proteins in Alzheimer’s disease pathology mouse models [13,15,57].

Overall, we show that within the microglial population, subsets of microglial cells shift to elongated, less ramified shapes that potentially shield the brain parenchyma from blood-antigens or mechanical stress one day after FUS-BBB modulation. This shift is in line with more frequent interactions between microglia and vessels showing astrocytic alterations, as well as increased microglial metabolic demand and disrupted digestive activity. In parallel, microglia reduced their frequency of interactions with pre- synaptic elements and extracellular space pockets. Therefore, within one day of FUS-BBB modulation, part of the microglial cell population may possibly focus on restoring BBB changes, such as swollen astrocytic endfeet, instead of pre-synaptic plasticity. This effect is not generalized to all the microglial population and warrants further investigation to clarify the long-term outcomes of FUS-BBB modulation on brain health.

### Limitations

This study has several limitations. First, increasing evidence demonstrate that microglia exhibit specific phenotypic responses in females compared to males, including increased inflammatory responses following BBB permeability in female mice . However, similar to the present study, previous research in the field used mainly male rodents [15]. Future studies investigating female cohorts and sex differences are highly encouraged, as they may prove beneficial in improving therapeutic FUS-BBB modulation for the female population. Second, though our analysis points to several microglial effects, we could not pinpoint the levels of influence of FUS + microbubbles, mechanical or otherwise, and blood-evading antigens and changes taking place through intermediaries, such as astrocytes, on the microglial responses. Astrocytes can instruct microglia to seal astrocytic endfeet gaps within the BBB in certain contexts, such as during postnatal development [62–64]. Also, several studies have provided evidence of astrocytic modulation following FUS-BBB [review 58]. Thus, assessing the dynamic crosstalk between microglia and astrocytes in the context of FUS-BBB modulation will further clarify the relevance of microglial mechanisms to the outcomes of FUS [66].

## Materials and Methods

### Animals and ethics

Male (n = 6 animals) C57BL/6J mice, 3 months of age, were purchased from the Jackson Laboratory and group-housed at 18-22 °C, 40-60% humidity and a 12 h light/dark cycle with *ad libitum* access to food and water. All the animal experiments were conducted following ethical approval from the Sunnybrook Research Institute Animal Care Committee and according to the Canadian Council on Animal Care Policies & Guidelines and the Animals for Research Act of Ontario.

### Magnetic resonance imaging-guided focused ultrasound sonication with microbubbles

Mice were anaesthetized using isoflurane at 2–3% with medical air as the gas carrier for 30 min (CP0406V2, Fresenius Kabi, Toronto, Canada). Each mouse was placed in a supine position on an MRI-compatible sled, with an angiocatheter inserted into the tail vein. The sled was inserted in a 7.0 T MRI (BioSpin 7030, Bruker, Massachusetts, United States of America [USA]) used to target the ventral hippocampi in the left hemisphere (ipsilateral) with a single focus beam, while the right hemisphere (contralateral) served as a control. The ultrasound waves were generated by a 0.58 MHz spherically focused transducer (75 mm outer diameter, 26 mm inner diameter, 60 mm radius of curvature) driven at the 3rd harmonic (1.78 MHz) and applied for 120 s in 0.01 s bursts at a frequency of 1 Hz, while 0.02 mL/kg Definity® MB (Lantheus Medical Imaging, Massachusetts, USA) were injected via the angiocatheter. The acoustic pressure was increased incrementally after each burst until subharmonic emissions were detected, when the acoustic pressure was reduced to 50% and maintained throughout the ultrasound treatment [67]. Subharmonic emissions were detected by a 16 mm diameter PZT hydrophone in the centre of the transducer and analyzed as described previously [68]. Immediately after, 0.2 mL/kg gadodiamide diluted in water (0.1 mmol/kg, 573.66 Da) MRI contrast agent (Gadovist administered at 0.2ml/kg, Bayer Inc., Mississauga, Ontario, Canada) was administered via the angiocatheter, followed by T1-weighted scans to visualize BBB permeability. The latter was quantified using Medical Image and Processing, Analysis and Visualization software (V11.0.17 for Mac OS X 10.7, MIPAV, National Institutes of Health, Maryland, USA) and regional-based analysis, as previously described [69]. Briefly, a 1 × 1 mm^2^ square, the theoretical size of the focused ultrasound spot, was used to measure the average voxel intensity in each ipsilateral targeted and corresponding non-targeted contralateral brain region [69].

### Perfusion and tissue sectioning

After 1 h and 24 h post FUS, male (n = 3 animals/time point) adult mice were anaesthetized with a mix of ketamine (80 mg/kg)/xylazine (10 mg/kg) and transcardially perfused with phosphate-buffered saline (PBS; 50 mM, pH 7.4), followed by 3.5% acrolein and 4% paraformaldehyde diluted in phosphate buffer (PB; 100 mM, pH 7.4). Brains were post-fixed in 4% paraformaldehyde diluted in PBS for 2 h at 4°C and the ipsilateral and contralateral brain hemispheres were separated. 50 μm longitudinal brain sections were next prepared in ice-cold PBS using a vibratome (VT1200S, Leica Biosystems, Ontario, Canada) at a frequency of 90–100 Hz and a speed of 0.5 mm/s. The sections were stored at -20°C in cryoprotectant (30% (v/v) glycerol and 30% (v/v) ethylene glycol in PBS) until further processing.

### Double immunohistochemistry and brightfield imaging

Double immunohistochemistry staining against Iba1 and IgG was performed in 3 sections containing the ventral hippocampi (Bregma -2.36 mm to -3.44 mm) from the ipsilateral and contralateral hemispheres per animal (n = 3 animals/time point) [70]. Sections were first assessed for IgG staining, used to delineate regions of increased BBB permeability after FUS-BBB modulation [13]. Briefly, free-floating sections were washed in PBS and quenched with 2% H_2_O_2_ diluted in 70% methanol for 10 min, followed by 0.1% NaBH_4_ diluted in PBS for 30 min. After 3 additional PBS washes, sections were incubated in a blocking solution of 10% normal donkey serum and 1% Triton X-100 in Tris-buffered saline (TBS; 50 mM, pH 7.4) for 1 h at room temperature (RT) and subsequently with donkey anti-mouse IgG secondary antibody (1:500 in blocking buffer, cat# 715-065-150, Jackson ImmunoResearch, Philadelphia, USA) for 2 h at RT. Following 3 washes in TBS, the sections were immersed in an avidin-biotin solution (1:100 in TBS, cat# VECTPK6100, VECTASTAIN, Vector Labs, California, USA) for 1 h at RT. The staining was next revealed by incubation with a HRP substrate kit solution (cat# SK-4600, Vector® VIP, Vector Labs, California, USA). The next day, the sections were stained for the marker Iba1, which is expressed by microglia in the brain parenchyma, as well as peripheral macrophages and border-associated macrophages [3]. It should be noted that the FUS-BBB modulation settings we utilized are not typically associated with peripheral immune cell infiltration into the brain [15], and only parenchymal cells were analyzed. We thus designated Iba1+ cells as microglia across the manuscript. Briefly, for Iba1 staining, sections were immersed in a blocking solution of 10% fetal bovine serum, 3% bovine serum albumin and 1% Triton X- 100 in TBS for 1 h at RT. Subsequently, the sections were incubated with a rabbit anti-Iba1 primary antibody (1:1000 in blocking buffer, cat# 019-19741, FUJIFILM Wako Chemical, Virginia, USA) overnight at 4°C. The following day, after 3 consecutive TBS washes, the sections were incubated in a biotinylated goat anti-rabbit secondary antibody (1:300 in TBS, cat# 111-066-046, Jackson Immunoresearch, Philadelphia, USA) and subsequently in an avidin–biotin solution for 1 h. Staining was revealed with 0.05% 3-3′-diaminobenzidine (DAB, D5905-50TAB, Millipore Sigma, Massachusetts, USA) and 0.015% H_2_O_2_ in 100 mM Tris–HCl. After mounting and drying, the sections were incubated in ultrapure H_2_O, increasing ethanol concentrations (50%, 70%, 80%, 90%, 100%) and xylene (cat# 534056, Millipore Sigma, Massachusetts, USA) for 5 min each at RT. The glass slides were cover-slipped with distyrene, plasticizer, and xylene mounting medium (cat# 13510, Electron Microscopy Sciences, Pennsylvania, USA). Each hemisphere was imaged (n = 3 animals/time point) in a single z plane at 40× with numerical aperture of 0.65, using a Basler area scan camera (2.3 MP; acA1920-40uc) and visualized through the Microvisioneer manualWSI Scan software (2019B-3S; Microvisioneer, Baden-Württemberg, Germany).

### IgG qualitative and quantitative brightfield analysis

All analyses were performed blinded to the experimental conditions using FIJI software (V2.13.1 for Mac OS X 10.7, FIJI, National Institutes of Health, Wisconsin, USA) [71], as previously described [43]. First, the freehand selection tool was used to trace the regions of interest (ROI), i.e. *CA1* and its strata, *Or*, *Py*, *Rad*, *LMol* [70]. Considering the structural diversity within these layers [56,74], each stratum was analyzed individually. Next, the IgG staining was visually identified in the ROI through Vector® VIP substrate deposition and measured in 3 sections (in both the ipsilateral and contralateral hemispheres per animal; n = 3 animals/time point), using an optical density analysis, as previously described [48].

### Microglial density and distribution analyses

The Iba1+ cell density and distribution in the *CA1* and its strata were assessed in 3 sections (in both the ipsilateral and contralateral hemispheres per animal; n = 3 animals/time point) via a semi-automated macro. In total, 3,079 Iba1+ cells (minimum of 227 microglia/layer/time point), were counted, a sample size which was considered sufficient to obtain statistical power based on calculations obtained using G*Power software (V3.1.9.6 for Mac OS X 10.7, G* Power Software, Nordrhein-Westfalen, Germany) (effect size of 0.25 and power of 0.8 estimated to 128 individual cells) [72]. Briefly, each Iba1+ cell in focus presenting a minimum of 3 processes was marked with the brush tool. Next, the ROIs were processed utilizing the grayscale, the thresholding and the plugins “analyze particles” as well as “nearest neighbour distance, NND”. Microglial density (# microglia/μm^2^) was defined as the total number of Iba1+ cells divided by the area. The NND (μm) was obtained by quantifying the average distance between each Iba1+ cell to its nearest neighbour. The spacing index (arbitrary unit, a.u.) was calculated as the square of the average NND multiplied by the microglia density. Two or more cells less than 12 µm apart were considered a cluster and averaged by animal to obtain the cluster density (# clusters/μm^2^) [73].

### Microglial morphology analysis

*LMol* Iba1+ cells were randomly selected in 3 sections (in both the ipsilateral or contralateral hemisphere) (n = 21 cells/hemisphere, N = 3 animals/time point) for the morphology analysis. This sample size was sufficient to obtain statistical power based on calculations using the G*Power software [72]. A previously developed semi-automated macro was adapted to this analysis [58]. Briefly, after tracing the cell body using the freehand selection tool, the following shape descriptors were obtained: area (μm^2^), perimeter (μm), circularity (4π × (area/perimeter^2^)), aspect ratio (major axis/minor axis, a.u.), roundness (arbor area/area of a circle with same convex perimeter, a.u.) and solidity (area/convex cell area, a.u.). A circularity of 1.0 represents a perfect circle and 0.0 is indicative of an elongated cell shape. Similarly, values higher than a 1.0 aspect ratio correspond to more elongated cell shapes. A solidity of 1.0 reflects a less ramified, convex shape, whereas 0.0 solidity points to a porous cell shape that is more ramified. Moreover, values closer to 1.0 for roundness represent more circular cell shapes [35].

Next, the polygon tool was used to trace the endpoints of the microglial processes, creating a convex shape that served as a proxy for the cellular territory, described through the aforementioned shape descriptors. The convex shape was subsequently processed to produce an unsharp mask of the cell, using the “remove outliers”, “clear outside” and “despeckle” functions. When necessary, the unsharp mask was manually corrected using the brush function to represent the observed cell. Once polished, the mask was used to obtain the preceding shape descriptors. The same mask was processed with the “analyze particles” and “analyze skeleton” commands to calculate the number of branches, endpoints, average branch length, maximum branch length (μm) and longest shortest path (μm), increasing with cell ramification. The mask arbor was also converted into an outline and analyzed using the FracLac [74] plugin to obtain the fractal dimension (a.u.) and lacunarity (a.u.), as previously described [35]. Fractal dimension and lacunarity are complementary measures [74], with higher values indicating a more complex organization of branching and more ramified morphological states [35]. Albeit informative, morphological analyses can be challenging to extrapolate across research teams [56] and represent one level of complexity of microglial function that should be complemented [7].

### Scanning electron microscopy staining, processing and imaging

Double immunohistochemistry staining against Iba1 and IgG was performed in 3 sections (in both the ipsilateral and contralateral hemispheres per animal; n = 3 animals/time point) using a similar protocol as described above. A few steps in the protocol were adapted to improve tissue preservation at the nanometric scale [19]. The quenching was performed with 0.3% H_2_O_2_ in PBS for 7 min; the blocking buffer and antibody incubation solutions contained 0.01% Triton X-100 and, after the DAB incubations, the sections were left in PB overnight at 4°C.

The following day, sections were post-fixed flat using an osmium-thiocarbohydrazide-osmium protocol [35], in which samples were first incubated in 3% potassium ferrocyanide (cat# PFC232.250, BioShop, Ontario, Canada) diluted in PB and combined (1:1) with 4% aqueous osmium tetroxide (cat# 19170, Electron Microscopy Sciences, Pennsylvania, USA) for 1 h at RT. After washes with ultrapure H_2_O, the sections were immersed in 1% thiocarbohydrazide (cat# 2231-57-4, Electron Microscopy Sciences, Pennsylvania, USA) for 20 min and an additional layer of 2% osmium tetroxide was applied for 30 min, with both solutions diluted in ultrapure H_2_O at RT. This was followed by dehydration with ascending concentrations of ethanol (35%, 50%, 70%, 80%, 90%, 100%) and propylene oxide (cat# 110205, Millipore Sigma, Massachusetts, USA) for 5 min each at RT. Next, sections were embedded in Durcupan ACM resin (cat# 44611-44614, Millipore Sigma, Massachusetts, USA) overnight at RT. The following day, sections were polymerized with a thin layer of resin between two fluoropolymer sheets (ACLAR; cat# 50425-25, Electron Microscopy Sciences, Pennsylvania, USA) placed for 72 h at 55 °C. The *CA1 LMol* was excised from the flat-embedded sections on ACLAR® sheets and glued to the top of resin blocks (2/hemisphere/animal, n = 3 animals/time point). Ultrathin sections (∼ 75 nm) were generated with an ultramicrotome (ARTOS 3D ultramicrotome, Leica Biosystems, Ontario, Canada), collected on a silicon nitride chip, and glued onto specimen mounts for SEM. In each resin block, 6 levels (∼ 5 μm apart) of ultrathin sections were collected. One ultrathin section was imaged per level using a Zeiss Crossbeam 350 Focused-Ion Beam SEM (Zeiss, Baden-Württemberg, Germany). Iba1+ microglial cell body were selected and the images exported as TIFF files with the Zeiss ATLAS Engine 5 software (Fibics, Ontario, Canada) at a resolution of 5 nm per pixel.

### Ultrastructural analyses

To quantify ultrastructural changes, we analyzed 151 microglial cell bodies (N = 11-13 microglial cell bodies from each ipsilateral and contralateral hemisphere, n = 3 animals/time point). This sample size was considered sufficient to obtain statistical power based on calculations using G*Power software [19,72]. Images were analyzed using QuPath V0.4.3 [78] and FIJI [73], adapted for use in previous work from our lab [44]. Microglial cell bodies were identified based on DAB staining, their smaller cell bodies and nuclei than neighbouring astrocytes or neurons, a characteristic heterochromatin pattern, long stretches of ER cisternae, as well as a presence of inclusions (e.g., lysosomes and lipofuscin granules) dispersed in their cytoplasm [19,35,36]. Microglial processes were identified based on their positive staining for Iba1 but were only analyzed qualitatively. Prior to analysis, a total of 3 trained and blinded observers separately agreed on the microglial identity of the selected cells.

Microglial cell body contacts with other cell bodies (i.e., astrocytes and neurons), myelinated axons, blood vessels, and synaptic elements (pre-synaptic axon terminals and post-synaptic spines) were quantified by an experimenter blinded to the experimental conditions [26,56,80]. The ultrastructural criteria used to identify each structure are outlined below. Astrocytes and astrocytic endfeet had pale nuclei with a thin rim of heterochromatin and pale irregular cytoplasm, often containing intermediate filaments [75]. Astrocytic endfeet were considered swollen when increased drastically in size, showing a clear cytoplasm enclosed by a plasma membrane [41]. Neurons presented pale nuclei and cytoplasm, and direct contacts with pre-synaptic terminals [75]. Myelinated axons were characterized by electron-dense sheaths and granular cytoplasm, often presenting mitochondria. Degraded myelin was recognized by ballooning, swelling or distancing between the well-defined myelin sheaths [75]. Blood vessel area (major radius*minor radius*π) was recorded using the freehand tool after two orthogonal diameters were traced, each delimited by the basement membrane, which was identified as an electron-dense layer surrounding endothelial cells forming blood vessels. The distance between each microglia membrane (from either a cell body or process) and the closest basement membrane was also traced [19,35,36].

Microglia were associated with a blood vessel when their distance from the vascular basement membrane was under 150 nm [19,35,36]. Moreover, among the parenchyma, microglial contacts with erythrocytes, which are recognized by their shape, size, lack of a nucleus and mitochondria, and completely electron-dense cytoplasm, were counted. Microglial contacts with synaptic elements were categorized as contacts with axon terminals or dendritic spines [19,35,36]. Pre-synaptic axon terminals showed a minimum of five synaptic vesicles, and were usually in contact with post-synaptic spines displaying a visible post- synaptic density [75]. Extracellular space pockets, essential for microglial motility and neuronal remodeling, were classified based on clear spaces without delineating membranes directly surrounding the microglia [43,44]. Extracellular digestion referred to extracellular space pockets containing debris in the vicinity of a microglial cell body or process, often in proximity to degraded myelin (ballooning, swelling or distancing between the well-defined myelin sheaths) [41].

The intracellular ultrastructural state of microglia, e.g., ER/Golgi apparatus, lysosomes, lipofuscin, mitochondria, and phagosomes, were also characterized [19,35,36]. ER/Golgi cisternae were identified by their long and narrow stretches. ER/Golgi apparatus dilation, associated with cellular stress, was identified when the distance between cisternal membranes was greater than 100 nm [17,19]. Mitochondria (homeostatic) were found in the cytoplasm, presenting an electron-dense appearance, double membrane, numerous cristae, and a circular shape. Mitochondria bigger than 1 μm in length were established as elongated, a phenomenon typically observed in response to cellular stress [76]. Moreover, when presenting with deteriorated outer membranes, vacuoles or degradation in their cristae membranes (electron-lucent pockets), mitochondria were classified as dystrophic [19,35,36]. Events of proximity between mitochondria and ER/Golgi apparatus cisternae, associated with calcium transfer and lipid synthesis [77], were quantified. Moreover, endothelial, astrocytic endfeet and synaptic element containing mitochondria were counted. Mitochondrial apposition to plasma membranes has been proposed to represent a component of purinergic signaling attracting microglial cell or process contacts [40]. Primary lysosomes were identified by their circular and homogenous contents (digestive enzymes) enclosed by a single membrane. Secondary lysosomes were darker, at least twice larger than primary lysosomes and often fused with phagosomes containing digested materials. Tertiary lysosomes were the largest, presenting residual materials, such as lipofuscin, large lipid bodies and phagosomes [19,35,36]. Lipid bodies are sites for synthesis and storage of immune mediators, including inflammatory cytokines [78], and were characterized by their electron-dense circular shape and interior. Oval structures with electron-dense content and a unique fingerprint-like pattern were identified as lipofuscin. Phagosomes were defined by their ovoid or circular shape with a single membrane and electron-lucent interior. They were classified as empty (completely electron-lucent) or filled (electron-lucent with content) [19,35,36].

Autophagosomes, part of an intracellular degradation pathway, were instead identified by the presence of elements inside circular, double-membrane vacuoles with a clear interior, resembling the cellular cytoplasm [19,79].

### Statistics

Microglial density, distribution, morphology and ultrastructural statistical analyses were conducted using the software GraphPad Prism (V 8.0.0 for MAC OS Ventura 13.4, GraphPad Software, California USA). Normality of the data was assessed using a Shapiro-Wilk test. A two-tailed paired Student’s t-test was used to compare the average contrast intensity between ipsilateral and contralateral *LMol*. Mixed effects 2-way ANOVA with Šídák’s multiple comparison correction was used for microglial density and distribution, optical density, morphology, and ultrastructure analyses comparing animal (n) averages between ipsilateral and contralateral *LMol* at the two examined time points [44]. As this is an exploratory study, all post-hoc pairwise comparisons were done, though only analyses comparing ipsilateral *vs* contralateral and 1 h *vs* 24 h are reported in the text. Both nonparametric and parametric data were used for the 2-way ANOVA, considering the lack of a non-parametric alternative to this test. The frequency distribution of cell values (morphology and ultrastructure) was assessed using the Gaussian or sum of 2 Gaussian nonlinear regression models for modal and bimodal data, respectively. The Wilcoxon test was applied to the nonlinear models comparing ipsilateral *vs* contralateral *LMol* in each time point [73]. When the number of recorded events within the cell dataset was not high enough to create the nonlinear model, we computed the presence *vs* absence of the variable across conditions. The conditional relative frequency of ultrastructural changes was measured using Fisher’s exact test. Spearman r correlations were computed for density and distribution, morphology and ultrastructural datasets separately, as described below. Mean differences were considered statistically significant when p < 0.05, with **** p < 0.0001, *** p < 0.001, ** p < 0.01, * p < 0.05. All reported values are presented as mean ± standard error of the mean (S.E.M).

#### Correlation analyses

The ipsilateral *CA1* density and distribution (n = 3 animals/time point) dataset and the optical density values (IgG staining) were imported into the GraphPad Prism built-in correlation tool to compute relationships between measured features. The same approach was used to explore the ipsilateral relationships between the 30 morphological features analyzed and optical density (n = 3 animals/time point) (Table S5). In addition, contralateral and ipsilateral correlations between the microglial cell body distance to the closest vessel and vessel area to all microglial ultrastructural features were analyzed (n = 151 cells, N = 3 animals/time point) (Table S11).

#### Principal component analyses

The Graph-Pad Prism built-in component analysis (PCA) was used to reduce dimensionality of the ipsilateral and contralateral morphology and ultrastructure datasets. Briefly, the variables were standardized to have a mean 0 and standard deviation of 1 to obtain a correlation matrix. The PC were selected based on a parallel analysis. Moreover, loadings and PC scores were visualized using scatterplots. Loadings and PC scores were used to understand the morphological and ultrastructural features that contributed to most of the variability in the ipsilateral compared to the contralateral *LMol* datasets.

## Acknowledgements

The authors would like to thank M.Sc. Kristina Mikloska for aiding in the operation of the FUS and MRI equipment, and Dr. Haley A. Vecchiarelli for the guidance provided during the project. We acknowledge and respect that the University of Victoria is located on the territory of the ləkwəŋən peoples and that the Songhees, Esquimalt, and WSÁNEĆ peoples have relationships to this land.

## Author contributions

This study was designed by E.G.A, R.H.K, I.A., and M.E.T. The manuscript was written and the figures were prepared by E.G.A, with supervision of I.A., and M.E.T. MRI-guided FUS was carried by R.H.K. with supervision of I.A. and K.H. Perfusions were performed by K.P. Light and electron microscopy staining were performed by K.H.P. and E.G.A. Light microscopy imaging were performed by K.H.P. and E.G.A. Electron microscopy imaging was performed by J.V., M.K. and F.G.I. Density analysis was performed by K.H.P. Optical density, morphology and ultrastructural analyses were performed by E.G.A. Experimental training was provided by K.P., M.C., M.K., and F.G.I. Overall supervision and support were provided by I.A. and M.E.T.

## Data and materials availability

All data generated during this study are included in this manuscript and its supplementary information files. The datasets analysed in the current study can be made available from the corresponding authors upon reasonable request.

