## Supplementary Files for "Early micro and nanoscopic responses of microglia to blood-brain barrier modulation by transcranial-focused ultrasound"

Microglia after transcranial-focused ultrasound

*Co-correspondence:

Marie-Ève Tremblay:

Isabelle Aubert:

### Supplementary Materials

#### Supplementary Methods

The nonlinear regression models were characterized based on their reported amplitude and mean (Table S5, 10). A higher curve amplitude indicates a narrower distribution of values, therefore, less variability in that morphological feature across cells. A higher curve mean reflects a shift toward larger values of the feature across the population. We observed microglia in the ipsilateral hemisphere deviated to larger and rounder somas with porous shapes, based on higher mean area, perimeter, roundness, but lower solidity at 1 h and 24 h following FUS-BBB (Figure 3K-P). In parallel, at 1 h there was a more homogenous distribution of microglial soma area and compactness, based on the higher amplitude of soma area, roundness, solidity, and aspect ratio; although the inverse was true for soma perimeter and circularity amplitudes, which were lower, therefore, there was a larger spread of the distribution. At 24 h, the relative distribution of all soma shape descriptors became more heterogeneous, that is, there was lower amplitude in the ipsilateral compared to the contralateral *LMol,* except for the aspect ratio (Figure 3K-P).

Regarding the processes, microglial cells appeared to adopt smaller, given the lower mean area and perimeter, and more regular process shapes, as per the lower mean roundness, solidity, lacunarity, fractal dimension, of the regression curves in the ipsilateral *LMol* at 1 h. At 24 h, the process masks shifted back to larger, indicated by a higher mean area, more compact, as per a lower mean perimeter and roundness, and irregular shapes, due to higher mean solidity, lacunarity and fractal dimension, in the ipsilateral compared to *LMol* regressions (Table S5). Correspondingly, there was a more heterogeneous relative distribution of all process descriptors at 1 h, that is, lower amplitude, except for the area in the ipsilateral *vs* contralateral *LMol* (Table S5). The opposite was observed at 24 h, whereby there was a homogenous relative distribution of process mask size, indicated by higher area and perimeter amplitude, and shape complexity, referred to by higher fractal dimension and roundness amplitude; although porosity, represented by lower solidity and lacunarity amplitudes, remained heterogeneous (Table S5).

In the ultrastructural analysis, comparison of means of the nonlinear regression models suggested that the relative distribution of microglial contacts with pre-synaptic elements reduced at both time points, indicated by lower means in the ipsilateral compared to the contralateral *LMol*, resulting in a more homogeneous distribution of contacts at 1 h and the reverse effect at 24 h, represented by higher and lower amplitudes, respectively (Table S10). Moreover, at 1 h in the ipsilateral *LMol,* the relative distribution of microglial cell bodies with higher metabolic demands increased, identified by elevated ER/Golgi and elongated mitochondria means, but decreased homeostatic mitochondria means, compared to the contralateral regressions. At 24 h, microglial ER/Golgi means were still higher, but microglial mitochondria and elongated mitochondria means decreased in the ipsilateral compared to the contralateral regressions. Such shifts resulted in an increasingly ipsilateral heterogeneous ER/Golgi distribution across microglial cell bodies at 1 h and 24 h, revealed by higher and lower amplitudes, respectively, with the reverse effect observed for homeostatic and elongated mitochondria, which showed lower amplitudes (Table S10).

#### Supplementary Figures

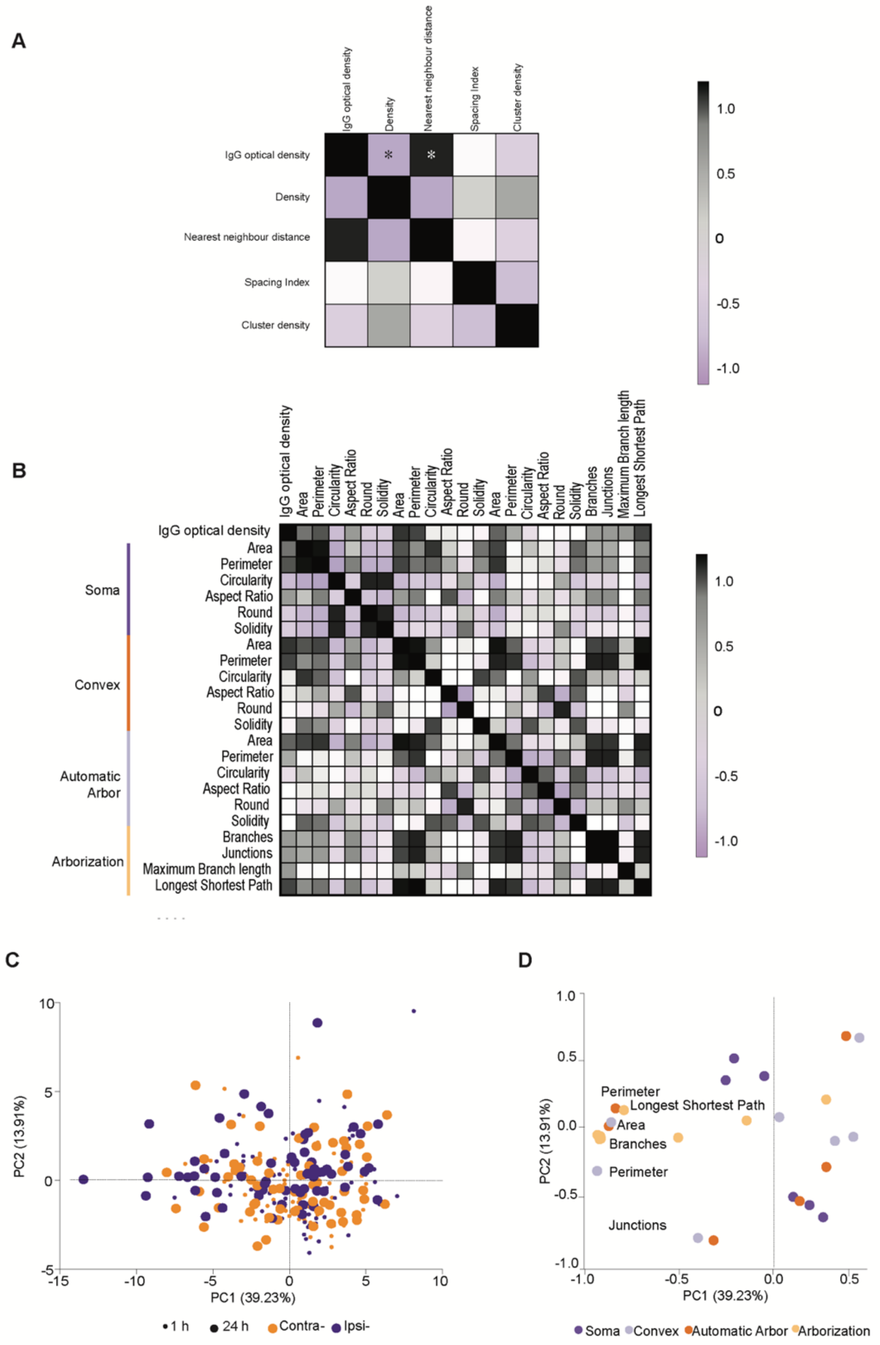


**Figure S1 Changes in microglial density and distribution correlated with IgG immunostaining intensity at 1 h and 24 h FUS-BBB modulation. A.** Optical density was significantly and negatively correlated with microglial density, with the reverse relationship observed for microglial nearest neighbour distance (NND) in the ipsilateral (ipsi-) *lacunosum moleculare* (*LMol*) at 1 hour (h) and 24 h following FUS-BBB modulation. **B.** Optical density did not correlate with any ipsilateral adaptations in morphology. Correlation matrices of ipsilateral features analyzed by Spearman r. **C.** Scores for the first and second principal component (PC1/PC2) of morphological features for all the cells analyzed, colored according to the *LMol* hemisphere (orange: contralateral, purple: ipsilateral) and time point (small icon: 1 h, big icon: 24 h). A higher variability in parameters is present at 24 h, particularly in the ipsilateral region group. **D.** Loading plot of morphology dataset, depicting the correlation between the morphological features (light orange: arborization, dark orange: convex shape, light purple: automatic arbor, dark purple: soma) and PC1/PC2, suggesting that variability in arborization descriptors contribute to overall differences between ipsilateral and contralateral *LMol* ionized calcium-binding adapter molecule 1 (Iba1) positive (+) cells.

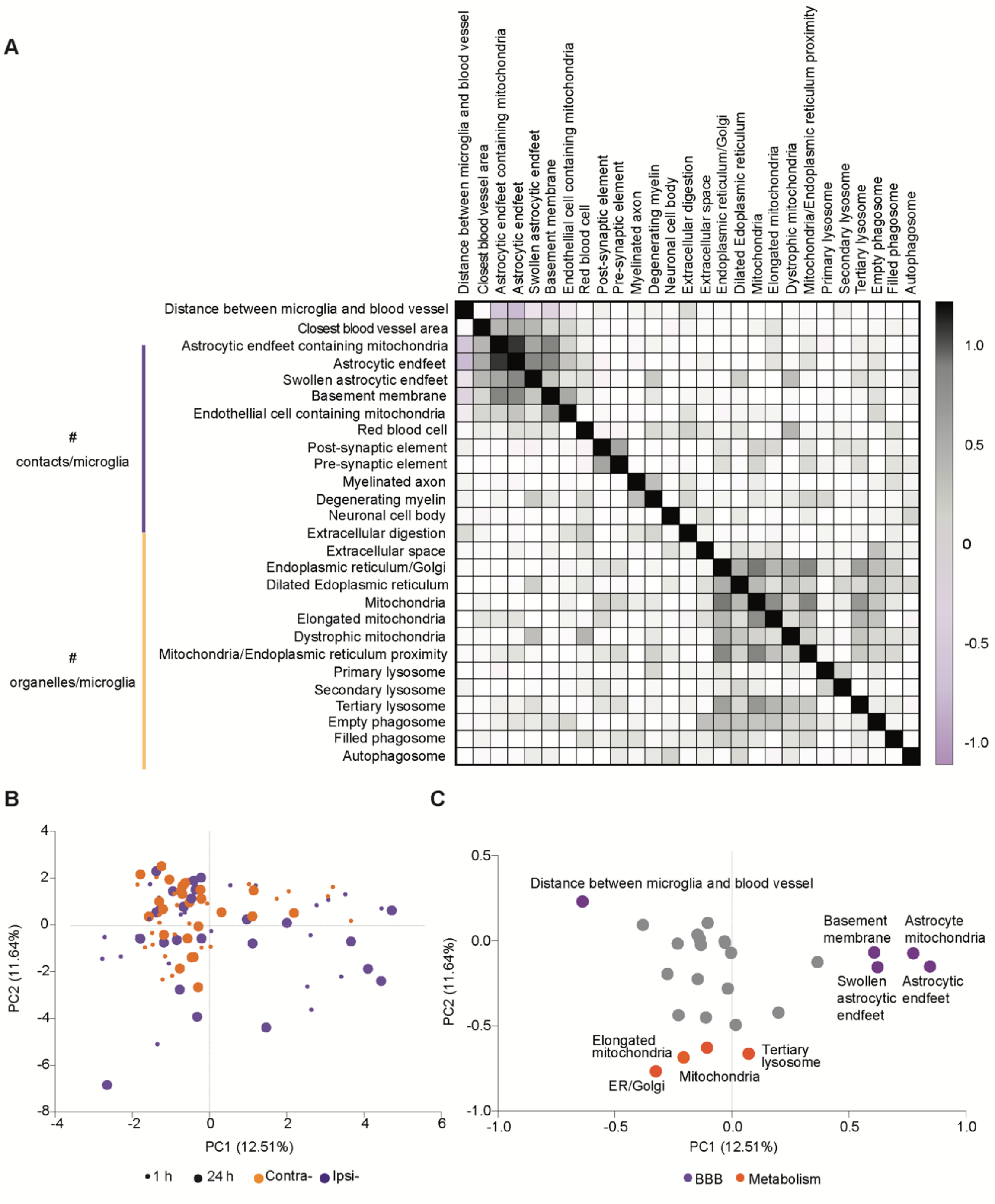
A
Distance between microglia and blood vessel
Closest blood vessel area
Astrocytic endfeet containing mitochondria
Astrocytic endfeet
Swollen astrocytic endfeet
Basement membrane
Endothellial cell containing mitochondria
Red blood cell
Post-synaptic element
Pre-synaptic element
Myelinated axon
Degenerating myelin
Neuronal cell body
Extracellular digestion
Extracellular space
Endoplasmic reticulum/Golgi
Dilated Edoplasmic reticulum
Mitochondria
Elongated mitochondria
Dystrophic mitochondria
Mitochondria/Endoplasmic reticulum proximity
Primary lysosome
Secondary lysosome
Tertiary lysosome
Empty phagosome
Filled phagosome
Autophagosome
### contacts/microglia
### organelles/microglia
1.0
0.5
0
-0.5
-1.0
B
PC1 (12.51%)
PC2 (11.64%)
1 h
24 h
Contra-
Ipsi-
C
Distance between microglia and blood vessel
Basement membrane
Astrocyte mitochondria
Swollen astrocytic endfeet
Astrocytic endfeet
Elongated mitochondria
Tertiary lysosome
Mitochondria
ER/Golgi
PC1 (12.51%)
PC2 (11.64%)
BBB
Metabolism

**Figure S2 The microglial ultrastructure varied with FUS-BBB modulation. A**. The microglial cell body distance from blood vessels was negatively correlated, while was positively correlated with changes in microglial cell body contacts with astrocytic endfeet, swollen astrocytic endfeet, basement membrane and endothelial cell containing mitochondria in the ipsilateral (ipsi-) *stratum lacunosum moleculare* (*LMol)* at 1 hour (h) and 24 h following FUS-BBB modulation. Correlation matrix of ipsilateral features analyzed by Spearman r. **B**. Scores for the first and second principal components 1 and 2 (PC1/PC2) for the microglial of ultrastructural features analyzed, colored according to the *LMol* hemisphere (orange: contralateral, purple: ipsilateral) and time point (small icon: 1 h, big icon: 24 h). A higher variability in parameters is present at 24 h, particularly in the ipsilateral *LMol*. **C**. Loading plot of ultrastructural dataset, depicting the correlation between the ultrastructural features (orange: microglial cellular stress, purple: microglial contacts with the blood-brain barrier or BBB) and PC1/PC2, suggesting that most adaptations between the ipsilateral and contralateral *LMol* ionized calcium-binding adapter molecule 1 (Iba1) positive (+) cells in the microglial ultrastructure scanning electron microscopy dataset arise from microglial cell body interactions with the BBB and cellular stress.

### Supplementary Tables

#### Table S1. Increased BBB permeability descriptive statistics.

*p* values are given after paired t-test of contra- and ipsi- average contrast intensity (Average contrast intensity) and 2-way ANOVA with Šídák's multiple comparisons test of optical density absolute values in each *CA1* strata (Optical density). For each data, SW results, mean and S.E.M of n = 3 animals/timepoint are given. ANOVA: analysis of variance, BBB: blood-brain barrier, Contra- or C: contralateral, Ipsi- or I: ipsilateral, SW: Shapiro-Wilk normality test, N: no, Y: yes, *p: p* value, f: f value, df: degrees of freedom and S.E.M. standard error of the mean.

| Average contrast intensity | Paired t test | | Contra- | | | Ipsi- | | |
|---|---|---|---|---|---|---|---|---|
| | df=5 | *p* | SW | Mean | S.E.M | SW | Mean | S.E.M |
| | t=2.66 | 0.044 | Y | 6629.0 | 981.4 | Y | 9421 | 1302 |

Optical density 2-way ANOVA

| df = 30 | *p* | f |
|---|---|---|
| Layer Time | 0.94 | 0.43 |
| Layer | 0.77 | 0.45 |
| Time | <0.0001 | 17.49 |
| Subject | 0.39 | 1.10 |

Optical Density - Šídák's multiple comparisons

| Comparison | *p* |
|---|---|
| C1xI1 | 0.8933 |
| C1xC24 | 0.9993 |
| **C1xI24** | **<0.0001** |
| I1xC24 | 0.9355 |
| **I1xI24** | **<0.0001** |
| **C24xI24** | **<0.0001** |

#### Table S2. Microglial density and distribution descriptive statistics.

*p* values are given after mixed effects 2-way ANOVA with Šídák's multiple comparisons test comparing averages per animal for each *CA1* strata across the two timepoints (1 h and 24 h). For each data, SW results, mean and S.E.M of n = 3 animals/timepoint/hemisphere are given. ANOVA: analysis of variance, BBB: blood-brain barrier, *CA1*: *cornus ammonis* 1, *LMol*: *stratum lacunosum moleculare*, *Or*: *stratum oriens*, *Py*: *stratum pyramidal* and *Rad*: *stratum radiatum* C or Contra-: contra-, I or Ipsi-: ipsi-, SW: Shapiro-Wilk normality test, N: no, Y: yes, p: p value, f: f value, t: t distribution, df: degrees of freedom, T: timepoint, H: hemisphere, h: hours, S.E.M. standard error of the mean and NND: nearest neighbour distance.

| | Contra- | | | Ipsi- | | |
|---|---|---|---|---|---|---|
| | SW | Mean | S.E.M | SW | Mean | S.E.M |
| | | Density (cells/µm$^2$) | | | | |
| *LMol* | Y | 0.00010 | 0.00000 | Y | 0.00010 | 0.00000 |
| *Rad* | | 0.00010 | 0.00000 | | 0.00020 | 0.00000 |
| *Py* | | 0.00010 | 0.00000 | | 0.00020 | 0.00000 |
| *Or* | | 0.00010 | 0.00000 | | 0.00010 | 0.00000 |
| *CA1* | | 0.00010 | 0.00000 | | 0.00020 | 0.00000 |
| | | Cluster density (cells/µm$^2$) | | | | |
| *LMol* | N | 0.00 | 0.00 | N | 0.00 | 0.00 |
| *CA1* | | 0.00 | 0.00 | Y | 0.00 | 0.00 |
| | | NND (a.u.) | | | | |
| *LMol* | Y | 53.20 | 2.20 | Y | 53.50 | 2.57 |
| *Rad* | | 53.30 | 1.41 | | 50.40 | 2.45 |
| *Py* | | 73.70 | 2.54 | | 69.00 | 0.68 |
| *Or* | | 56.50 | 2.67 | | 54.90 | 4.21 |
| *CA1* | | 68.00 | 4.38 | | 61.90 | 8.59 |
| | | Spacing index (a.u.) | | | | |
| *LMol* | Y | 0.35 | 0.01 | Y | 0.37 | 0.01 |
| *Rad* | | 0.41 | 0.03 | | 0.39 | 0.01 |
| *Py* | | 0.53 | 0.03 | | 0.49 | 0.08 |
| *Or* | | 0.40 | 0.02 | | 0.45 | 0.04 |
| *CA1* | | 0.76 | 0.06 | | 0.64 | 0.12 |

| Mixed effects 2-way ANOVA | | | |
|---|---|---|---|
| Layer | Main effect | p | f |
| | Density (µm2) | | |
| *LMol* | T | 0.13 | 3.6 |
| | H | 0.22 | 2.1 |
| | TxH | 0.35 | 1.1 |
| *Rad* | T | 0.28 | 1.3 |
| | H | 0.28 | 1.3 |
| | TxH | 0.25 | 1.6 |
| *Py* | T | 0.35 | 1.0 |
| | H | 0.67 | 0.2 |
| | TxH | 0.89 | 0.0 |
| *Or* | T | 0.22 | 2.1 |
| | H | 0.78 | 0.1 |
| | TxH | 0.42 | 0.8 |
| *CA1* | T | 0.16 | 2.4 |
| | H | 0.66 | 0.2 |
| | TxH | 0.95 | 0.0 |
| | Cluster density (µm2) | | |
| *LMol* | T | 0.50 | 0.5 |
| | H | 0.13 | 2.9 |
| | TxH | 0.13 | 2.9 |
| *CA1* | T | 0.82 | 0.1 |
| | H | 0.19 | 2.1 |
| | TxH | 0.36 | 0.9 |
| | NND (a.u.) | | |
| *LMol* | T | 0.71 | 0.2 |
| | H | 0.13 | 3.6 |
| | TxH | *0.06* | 6.8 |
| *Rad* | T | 0.18 | 2.7 |
| | H | 0.47 | 0.6 |
| | TxH | 0.49 | 0.6 |
| *Py* | T | 0.12 | 3.0 |
| | H | 0.44 | 0.7 |
| | TxH | 0.59 | 0.3 |

| Mixed effects 2-way ANOVA | | | |
|---|---|---|---|
| Layer | Main effect | p | f |
| *Or* | T | 0.81 | 0.1 |
| | H | 0.72 | 0.1 |
| | TxH | 0.89 | 0.0 |
| ***CA1*** | **T** | **0.05** | **5.5** |
| | H | 0.89 | 0.0 |
| | TxH | 0.86 | 0.0 |
| *LMol* | T | 0.31 | 1.3 |
| | H | 0.58 | 0.4 |
| | TxH | 0.43 | 0.8 |
| *Rad* | T | 0.18 | 2.6 |
| | H | 0.10 | 4.7 |
| | TxH | 0.55 | 0.4 |
| *Py* | T | 0.61 | 2.6 |
| | H | 0.67 | 4.7 |
| | TxH | 0.32 | 0.4 |
| *Or* | T | 0.39 | 0.8 |
| | H | 0.53 | 0.4 |
| | TxH | 0.76 | 0.1 |
| *CA1* | T | 0.07 | 4.2 |
| | H | 0.19 | 2.1 |
| | TxH | 0.87 | 0.0 |

**Table S3. Microglia density and distribution correlation with BBB permeability descriptive statistics.**

*p* values are given after Spearman r correlation between contrast enhancement and density, NND, spacing index, cluster density and optical density in the ipsi- *LMol*. For each data, SW results, mean, S.E.M, r and p of n = 3 animals/timepoint/hemisphere are given. NND: nearest neighbour distance, SW: Shapiro-Wilk normality test, N: no, Y: yes, p: p value, r: r coefficient and S.E.M. standard error of the mean.

| | | | | Density | | NND | | Spacing Index | | Cluster density | |
|---|---|---|---|---|---|---|---|---|---|---|---|
| | SW | Mean | S.E.M | r | p | r | p | r | p | r | p |
| Density | | 0.00 | 0.00 | 1.00 | | -0.94 | 0.02 | 0.20 | 0.71 | 0.38 | 0.47 |
| NND | | 53.46 | 1.41 | -0.94 | 0.02 | 1.00 | | -0.14 | 0.80 | -0.49 | 0.33 |
| Spacing Index | Y | 0.37 | 0.01 | 0.20 | 0.71 | -0.14 | 0.80 | 1.00 | | -0.70 | 0.14 |
| Cluster Density | | 0.00 | 0.00 | 0.38 | 0.47 | -0.49 | 0.33 | -0.70 | 0.14 | 1.00 | |
| Optical Density | | 1.17 | 0.07 | -0.94 | 0.02 | 0.89 | 0.03 | -0.09 | 0.92 | -0.52 | 0.30 |

#### Table S4. Microglial morphology descriptive statistics.

*p* values are given after a mixed-effects 2-way ANOVA with Šídák's multiple comparisons test comparing averages per animal across the ipsi- and contra- *LMol* across the two timepoints (1 and 24 h). For each data, SW results, mean and S.E.M of n = 3 animals/timepoint are given. ANOVA: analysis of variance, a.u.: arbitrary unit, #: number, *LMol*: *stratum lacunosum moleculare*, C or Contra-: contra-, I or Ipsi-: ipsi-, SW: Shapiro-Wilk normality test, *: N too small for normality assessment, p: p value, f: f value, t: t distribution, df: degrees of freedom, T: timepoint, H: hemisphere, h: hours and S.E.M. standard error of the mean.

| | Contra- | | Ipsi- | | Mixed effects 2-way ANOVA | | |
|---|---|---|---|---|---|---|---|
| Parameter | Mean | S.E.M | Mean | S.E.M | | f | p |
| | | | | Soma | | | |
| Area ($\mu m^2$) | 43.31 | 3.79 | 63.72 | 20.89 | T | 3.53 | 0.13 |
| | | | | | H | 3.35 | 0.14 |
| | | | | | TxH | 2.35 | 0.20 |
| Aspect Ratio (a.u.) | 1.88 | 0.11 | 1.93 | 0.08 | T | 4.76 | 0.06 |
| | | | | | H | 0.30 | 0.60 |
| | | | | | TxH | 0.11 | 0.75 |
| **Perimeter (a.u.)** | 29.46 | 0.13 | 39.58 | 7.15 | T | 1.82 | 0.25 |
| | | | | | **H** | **15.43** | **0.02** |
| | | | | | **TxH** | **8.01** | **0.05** |
| **Roundness (a.u.)** | 0.58 | 0.03 | 0.58 | 0.02 | **T** | **11.17** | **0.01** |
| | | | | | H | 0.11 | 0.75 |
| | | | | | TxH | 0.48 | 0.51 |
| Circularity (a.u.) | 0.57 | 0.02 | 0.54 | 0.05 | T | 3.06 | 0.16 |
| | | | | | H | 4.63 | 0.10 |
| | | | | | TxH | 4.63 | w0.10 |
| Solidity (a.u.) | 0.82 | 0.00 | 0.80 | 0.01 | T | 0.39 | 0.56 |
| | | | | | H | 1.68 | 0.26 |
| | | | | | TxH | 0.57 | 0.49 |
| | | | | Convex | | | |
| Area ($\mu m^2$) | 1084.00 | 24.79 | 1138.00 | 151.90 | T | 1.86 | 0.21 |
| | | | | | H | 0.18 | 0.69 |
| | | | | | TxH | 0.96 | 0.36 |
| Aspect Ratio (a.u.) | 1.69 | 0.01 | 1.67 | 0.01 | T | 0.00 | 0.99 |
| | | | | | H | 0.07 | 0.81 |
| | | | | | TxH | 0.07 | 0.81 |
| Perimeter (a.u.) | 144.20 | 1.13 | 145.80 | 6.65 | T | 0.34 | 0.58 |
| | | | | | H | 0.03 | 0.87 |
| | | | | | TxH | 0.67 | 0.44 |
| Roundness (a.u.) | 0.67 | 0.01 | 0.65 | 0.02 | T | 1.08 | 0.36 |
| | | | | | H | 1.48 | 0.29 |
| | | | | | TxH | 4.11 | 0.11 |
| Circularity (a.u.) | 0.64 | 0.01 | 0.64 | 0.00 | T | 0.07 | 0.79 |
| | | | | | H | 0.02 | 0.88 |
| | | | | | TxH | 0.01 | 0.91 |
| Solidity | 0.89 | 0.01 | 0.90 | 0.01 | T | 4.48 | 0.07 |

| | | | | | | | |
|---|---|---|---|---|---|---|---|
| (a.u.) | | | | | H | 1.62 | 0.24 |
| | | | | | TxH | 0.02 | 0.88 |
| **Automated cell mask** | | | | | | | |
| | | | | | T | 3.76 | 0.12 |
| Area ($\mu m^2$) | 261.60 | 6.73 | 288.50 | 46.17 | H | 1.17 | 0.34 |
| | | | | | TxH | 2.52 | 0.19 |
| | | | | | T | 0.01 | 0.91 |
| Aspect Ratio (a.u.) | 1.84 | 0.04 | 1.90 | 0.03 | H | 0.14 | 0.72 |
| | | | | | TxH | 0.22 | 0.65 |
| | | | | | T | 0.00 | 0.99 |
| Roundness (a.u.) | 0.60 | 0.00 | 0.59 | 0.00 | H | 0.22 | 0.65 |
| | | | | | TxH | 0.01 | 0.92 |
| | | | | | T | 0.00 | 0.99 |
| Fractal dimension (a.u.) | 1.39 | 0.01 | 1.38 | 0.01 | H | 0.05 | 0.83 |
| | | | | | TxH | 0.56 | 0.48 |
| | | | | | T | 0.76 | 0.41 |
| Lacunarity (a.u.) | 0.25 | 0.00 | 0.27 | 0.01 | H | 1.56 | 0.25 |
| | | | | | TxH | 0.17 | 0.69 |
| | | | | | T | 0.63 | 0.45 |
| Circularity (a.u.) | 0.03 | 0.01 | 0.03 | 0.00 | H | 0.01 | 0.91 |
| | | | | | TxH | 0.47 | 0.51 |
| | | | | | T | 0.31 | 0.59 |
| Perimeter (µm) | 388.60 | 3.95 | 386.20 | 29.25 | H | 0.00 | 0.96 |
| | | | | | TxH | 0.53 | 0.49 |
| | | | | | T | 0.76 | 0.41 |
| Solidity (a.u.) | 0.25 | 0.00 | 0.27 | 0.01 | H | 1.56 | 0.25 |
| | | | | | TxH | 0.17 | 0.69 |
| **Arborization** | | | | | | | |
| | | | | | T | 0.30 | 0.61 |
| Branches (#) | 61.81 | 0.84 | 63.78 | 5.29 | H | 0.08 | 0.79 |
| | | | | | TxH | 0.78 | 0.43 |
| | | | | | T | 0.30 | 0.61 |
| Junctions (#) | 30.33 | 0.34 | 31.49 | 2.60 | H | 0.11 | 0.76 |
| | | | | | TxH | 0.68 | 0.46 |
| | | | | | T | 0.39 | 0.55 |
| Longest Shortest Path (µm) | 69.91 | 1.09 | 71.80 | 3.72 | H | 0.20 | 0.67 |
| | | | | | TxH | 1.30 | 0.29 |
| | | | | | T | 0.20 | 0.66 |
| Max branch length (µm) | 14.43 | 0.10 | 15.32 | 0.19 | H | 1.88 | 0.21 |
| | | | | | TxH | 0.02 | 0.90 |

| Šídák's multiple comparisons test | | | | | | | | |
|---|---|---|---|---|---|---|---|---|
| | 1 h *vs* 24 h | | | | Contra- *vs* Ipsi- | | | |
| Parameter | H | p | t | df | T | p | t | df |
| **Soma** | | | | | | | | |
| Perimeter (a.u.) | C | 1.00 | 0.05 | | 1 | 0.73 | 0.78 | 4 |
| | I | 0.08 | 2.47 | | 24 | **0.02** | 4.78 | |

#### Table S5. Morphological Nonlinear regression parameters.

*p* values are given after paired Wilcoxon test of Contra- *vs* Ipsi- nonlinear regression curve fits. These were modeled after the relative distribution of each parameter in the two *LMol* (contra- and ipsi-) nested into the two timepoints (1 and 24 h). For each data, Amplitude, Mean and p value of n = 3 animals/timepoint/hemisphere are given. a.u.: arbitrary unit, #: number, p: p value, *LMol*: *stratum lacunosum moleculare*, Contra-: contra-, Ipsi-: ipsi- and h: hours.

| Nonlinear regression | | 1 h | | 24 h | |
|---|---|---|---|---|---|
| | | Contra- | Ipsi- | Contra- | Ipsi- |
| Soma | | | | | |
| Area (µm²) | Amplitude | 60.20 | 62.57 | 36.40 | 20.91 |
| | Mean | 37.09 | 38.20 | 34.36 | 59.99 |
| | p | <0.0001 | | <0.0001 | |
| Aspect Ratio (a.u.) | Amplitude | 47.24 | 58.09 | 32.00 | 37.24 |
| | Mean | 1.55 | 1.55 | 1.86 | 1.72 |
| | p | <0.0001 | | <0.0001 | |
| Circularity (a.u.) | Amplitude | 13.82 | 13.59 | 15.02 | 16.62 |
| | Mean | 0.59 | 0.57 | 0.57 | 0.49 |
| | p | <0.0001 | | 0.58 | |
| Perimeter (µm) | Amplitude | 61.38 | 46.52 | 47.87 | 28.05 |
| | Mean | 28.17 | 28.36 | 24.25 | 40.68 |
| | p | <0.0001 | | <0.0001 | |
| Roundness (a.u.) | Amplitude | 13.17 | 14.80 | 12.38 | 11.77 |
| | Mean | 0.62 | 0.63 | 0.53 | 0.55 |
| | p | <0.0001 | | <0.0001 | |
| Solidity (a.u.) | Amplitude | 21.81 | 23.14 | 27.40 | 23.78 |
| | Mean | 0.85 | 0.85 | 0.84 | 0.81 |
| | p | <0.0001 | | <0.0001 | |
| Convex arbor | | | | | |
| Area (µm²) | Amplitude | 18.97 | 17.54 | 17.89 | 16.03 |
| | Mean | 988.40 | 912.80 | 969.20 | 1083.00 |
| | p | <0.0001 | | 0.00 | |
| Aspect Ratio (a.u.) | Amplitude | 20.40 | 18.19 | 22.24 | 20.15 |
| | Mean | 1.49 | 1.51 | 1.42 | 1.52 |
| | p | <0.0001 | | <0.0001 | |
| Circularity (a.u.) | Amplitude | 14.45 | 14.76 | 17.06 | 17.90 |
| | Mean | 0.63 | 0.64 | 0.70 | 0.67 |
| | p | <0.0001 | | <0.0001 | |
| Perimeter (µm) | Amplitude | 18.97 | 20.54 | 21.53 | 21.59 |
| | Mean | 140.10 | 137.00 | 138.30 | 145.30 |
| | p | 0.00 | | 0.36 | |
| Roundness (a.u.) | Amplitude | 5.28 | 5.12 | 5.39 | 5.25 |
| | Mean | 0.18 | 0.18 | 0.20 | 0.17 |
| | p | <0.0001 | | <0.0001 | |
| Solidity (a.u.) | Amplitude | 11.46 | 9.73 | 10.92 | 13.77 |
| | Mean | 0.89 | 0.98 | 0.95 | 0.93 |

| | | | | | |
|---|---|---|---|---|---|
| | p | 0.73 | | 0.20 | |
| Automatic arbor | | | | | |
| Aspect ratio (a.u.) | Amplitude | 711.70 | 18.35 | 16.86 | 22.36 |
| | Mean | 1.65 | 1.47 | 1.49 | 1.61 |
| | p | <0.0001 | | <0.0001 | |
| Circularity (a.u.) | Amplitude | -33.45 | -325.9 | -45.64 | -16.89 |
| | Mean | 0.39 | 0.04 | 0.39 | 0.07 |
| | p | 0.0003 | | <0.0001 | |
| Perimeter (µm) | Amplitude | 15.09 | 13.70 | 10.51 | 15.08 |
| | Mean | 367.00 | 359.20 | 349.10 | 334.70 |
| | p | <0.0001 | | <0.0001 | |
| Roundness (a.u.) | Amplitude | 11.55 | 9.31 | 9.34 | 10.71 |
| | Mean | 0.63 | 0.60 | 0.62 | 0.58 |
| | p | 0.00 | | 0.00 | |
| Solidity (a.u.) | Amplitude | 20.36 | 17.41 | 22.63 | 16.83 |
| | Mean | 0.25 | 0.25 | 0.25 | 0.27 |
| | p | <0.0001 | | <0.0001 | |
| Area ($\mu m^2$) | Amplitude | 24.02 | 26.54 | 14.62 | 15.69 |
| | Mean | 241.80 | 187.30 | 223.00 | 231.80 |
| | p | <0.0001 | | 0.00 | |
| Fractal Dimension (a.u.) | Amplitude | 17.00 | 13.00 | 10.70 | 11.36 |
| | Mean | 1.40 | 1.38 | 1.38 | 1.39 |
| | p | 0.02 | | 0.00 | |
| Lacunarity (a.u.) | Amplitude | 26.83 | 19.28 | 22.77 | 17.35 |
| | Mean | 0.32 | 0.32 | 0.32 | 0.32 |
| | p | <0.0001 | | <0.0001 | |
| Arborization | | | | | |
| Branches (#) | Amplitude | 35.04 | 28.32 | 25.84 | 24.95 |
| | Mean | 56.75 | 57.23 | 59.33 | 53.89 |
| | p | <0.0001 | | <0.0001 | |
| Junctions (#) | Amplitude | 34.47 | 27.79 | 25.04 | 25.12 |
| | Mean | 28.04 | 27.86 | 29.37 | 26.37 |
| | p | <0.0001 | | <0.0001 | |
| Longest Shortest Path (µm) | Amplitude | 20.87 | 24.23 | 20.65 | 21.86 |
| | Mean | 70.54 | 69.25 | 64.74 | 71.46 |
| | p | <0.0001 | | <0.0001 | |
| Max Branch Length (µm) | Amplitude | 26.37 | 26.04 | 20.26 | 33.79 |
| | Mean | 13.65 | 13.54 | 13.07 | 12.60 |
| | p | <0.0001 | | <0.0001 | |

**Table S6. Morphology correlation with BBB permeability.**

*p* values are given after Spearman r correlation between optical density and morphology parameters in the ipsi- *LMol*. For each data, r and p of n = 3 animals/timepoint are given. p: p value, r: r coefficient.

| | Optical density | |
|---|---|---|
| Parameters | p | r |
| Soma Area | 0.24 | 0.60 |
| Soma Perimeter | 0.14 | 0.71 |
| Soma Circularity | 0.14 | -0.71 |
| Soma Aspect Ratio | 0.42 | 0.43 |
| Soma Roundness | 0.30 | -0.54 |
| Soma Solidity | 0.30 | -0.54 |
| Convex Area | 0.06 | 0.83 |
| Convex Perimeter | 0.10 | 0.77 |
| Convex Circularity | 0.92 | 0.09 |
| Convex Aspect Ratio | 0.92 | 0.09 |
| Convex Roundness | 0.92 | -0.09 |
| Convex Solidity | 0.80 | -0.14 |
| Automatic Arbor Area | 0.18 | 0.66 |
| Automatic Arbor Perimeter | 0.50 | 0.37 |
| Automatic Arbor Circularity | 0.50 | -0.37 |
| Automatic Arbor Aspect Ratio | 1.00 | -0.03 |
| Automatic Arbor Roundness | 0.92 | -0.09 |
| Automatic Arbor Solidity | 1.00 | -0.03 |
| Arborization Branches | 0.42 | 0.43 |
| Arborization Junctions | 0.42 | 0.43 |
| Maximum Branch length | 0.42 | 0.43 |
| Arborization Longest Shortest Path | 0.10 | 0.77 |

#### Table S7. Microglia morphology principal component analysis.

For each PC, eigenvalue, proportion of variance, cumulative proportion of variance and loadings for each morphology parameter of n = 21 cells, N = 3 animals/timepoint are given. PC: principal components.

| PC summary | PC1 | PC2 | PC3 | PC4 | PC5 | PC6 |
|---|---|---|---|---|---|---|
| Eigenvalue | 11.77 | 4.172 | 3.04 | 1.951 | 1.887 | 1.556 |
| Proportion of variance | 39.23% | 13.91% | 10.13% | 6.50% | 6.29% | 5.19% |
| Cumulative proportion of variance | 39.23% | 53.13% | 63.27% | 69.77% | 76.06% | 81.24% |
| Loadings | | | | | | |
| Soma Area | -0.29 | 0.44 | -0.49 | 0.18 | -0.02 | 0.09 |
| Soma Perimeter | -0.25 | 0.60 | -0.65 | 0.04 | -0.17 | 0.05 |
| Soma Circularity | 0.22 | -0.56 | 0.60 | 0.26 | 0.01 | -0.08 |
| Soma Aspect Ratio | -0.09 | 0.47 | -0.34 | -0.26 | 0.62 | -0.03 |
| Soma Round | 0.06 | -0.41 | 0.32 | 0.27 | -0.63 | 0.02 |
| Soma Solidity | 0.15 | -0.47 | 0.59 | 0.22 | 0.24 | -0.03 |
| Convex Area | -0.92 | 0.10 | 0.02 | 0.02 | -0.13 | -0.22 |
| Convex Perimeter | -0.88 | 0.23 | 0.21 | -0.15 | -0.15 | 0.04 |
| Convex Circularity | 0.10 | -0.44 | -0.48 | 0.34 | 0.18 | -0.58 |
| Convex Aspect Ratio | 0.35 | 0.76 | 0.44 | 0.23 | 0.02 | -0.08 |
| Convex Round | -0.36 | -0.73 | -0.44 | -0.27 | -0.07 | 0.13 |
| Convex Solidity | 0.24 | -0.19 | -0.42 | 0.46 | 0.08 | -0.66 |
| Automatic Arbor Area | -0.90 | 0.13 | -0.07 | 0.24 | -0.12 | 0.11 |
| Automatic Arbor Perimeter | -0.97 | 0.02 | 0.11 | -0.01 | -0.06 | -0.10 |
| Automatic Arbor Circularity | 0.38 | 0.03 | -0.20 | 0.46 | -0.20 | 0.47 |
| Automatic Arbor Aspect Ratio | 0.42 | 0.75 | 0.36 | 0.24 | 0.04 | -0.14 |
| Automatic Arbor Round | -0.44 | -0.71 | -0.35 | -0.27 | -0.11 | 0.19 |
| Automatic Arbor Solidity | 0.29 | 0.00 | -0.33 | 0.69 | -0.05 | 0.46 |
| Arborization Branches | -0.97 | 0.02 | 0.06 | 0.13 | 0.08 | 0.00 |
| Arborization Junctions | -0.96 | 0.02 | 0.06 | 0.13 | 0.08 | 0.00 |
| Maximum Branch length | -0.18 | 0.14 | -0.02 | -0.21 | -0.59 | -0.31 |
| Arborization Longest Shortest Path | -0.84 | 0.22 | 0.19 | 0.01 | -0.14 | -0.13 |
| Automatic Arbor Fractal Dimension | -0.76 | -0.22 | 0.07 | 0.20 | 0.13 | 0.09 |
| Automatic Arbor Lacunarity | -0.01 | 0.17 | 0.27 | -0.30 | 0.00 | 0.13 |

#### Table S8. Microglial ultrastructure descriptive statistics.

*p* values are given after a mixed-effects 2-way ANOVA with Šídák's multiple comparisons test comparing averages per animal across the ipsi- and contra- Lmol across the two timepoints (1 and 24 h). For each data, SW results, mean and S.E.M of n = 3 animals/timepoint are given. ANOVA: analysis of variance#: number, Lmol: *stratum lacunosum moleculare*, C or Contra- contra-, I or Ipsi-: ipsi-, SW: Shapiro-Wilk normality test, *: N too small for normality assessment, p: p value, f: f value, t: t distribution, df: degrees of freedom, T: timepoint, H: hemisphere, h: hours and S.E.M. standard error of the mean.

| | | Contra- | | Ipsi- | | Mixed effects 2-way ANOVA | | |
|---|---|---|---|---|---|---|---|---|
| Parameter | unit | Mean | S.E.M | Mean | S.E.M | | f | p |
| **BBB contacts** | | | | | | | | |
| Astrocytic endfeet containing mitochondria | #/cell | 0.17 | 0.12 | 0.24 | 0.04 | T | 0.65 | 0.45 |
| | | | | | | H | 0.55 | 0.48 |
| | | | | | | TxH | 2.48 | 0.15 |
| Astrocytic endfeet | | 0.24 | 0.13 | 0.51 | 0.08 | T | 1.36 | 0.31 |
| | | | | | | H | 2.67 | 0.18 |
| | | | | | | TxH | 0.11 | 0.76 |
| Swollen astrocytic endfeet | | 0.04 | 0.04 | 1.15 | 0.10 | T | 0.03 | 0.87 |
| | | | | | | **H** | **16.48** | **0.02** |
| | | | | | | TxH | 0.26 | 0.64 |
| Endothelial cell containing mitochondria | | 0.05 | 0.05 | 0.08 | 0.03 | T | 0.53 | 0.51 |
| | | | | | | H | 1.00 | 0.37 |
| | | | | | | TxH | 1.00 | 0.37 |
| Basement membrane | | 0.09 | 0.04 | 0.09 | 0.04 | T | 0.00 | 0.98 |
| | | | | | | H | 0.00 | 0.96 |
| | | | | | | **TxH** | **12.31** | **0.02** |
| Microglial blood vessel distance | μm | 6.82 | 1.20 | 7.34 | 0.13 | T | 0.36 | 0.56 |
| | | | | | | H | 0.09 | 0.77 |
| | | | | | | TxH | 0.56 | 0.47 |
| Density of microglia associated with vessels | #/μm$^2$ | 0.17 | 0.07 | 0.24 | 0.02 | T | 0.97 | 0.38 |
| | | | | | | H | 1.09 | 0.36 |
| | | | | | | TxH | 0.62 | 0.47 |
| Red blood cells contacts | #/cell | 0.00 | 0.00 | 0.05 | 0.05 | T | 1.00 | 0.37 |
| | | | | | | H | 1.00 | 0.37 |
| | | | | | | TxH | 1.00 | 0.37 |
| **Synaptic plasticity contact** | | | | | | | | |
| Synapse containing mitochondria contact | #/cell | 0.78 | 0.44 | 0.79 | 0.35 | T | 9.68 | 0.04 |
| | | | | | | H | 0.01 | 0.94 |
| | | | | | | TxH | 0.24 | 0.65 |
| Post-synaptic element | | 0.84 | 0.17 | 0.97 | 0.30 | T | 3.41 | 0.14 |
| | | | | | | H | 0.35 | 0.59 |
| | | | | | | TxH | 0.35 | 0.59 |

| | | | | | | | | |
|---|---|---|---|---|---|---|---|---|
| Pre-synaptic element | | 2.82 | 1.11 | 2.74 | 1.02 | **T** | **33.06** | **0.00** |
| | | | | | | H | 0.05 | 0.82 |
| | | | | | | TxH | 0.05 | 0.82 |
| Satellite | | 0.09 | 0.01 | 0.11 | 0.01 | T | 0.01 | 0.91 |
| | | | | | | H | 0.09 | 0.77 |
| | | | | | | TxH | 0.09 | 0.77 |
| Myelinated axon | | 0.13 | 0.03 | 0.20 | 0.06 | T | 1.16 | 0.31 |
| | | | | | | H | 0.74 | 0.41 |
| | | | | | | TxH | 0.19 | 0.68 |
| Degenerating myelin | | 0.07 | 0.01 | 0.08 | 0.03 | T | 0.56 | 0.47 |
| | | | | | | H | 0.06 | 0.82 |
| | | | | | | TxH | 0.08 | 0.79 |
| Extracellular digestion | | 0.17 | 0.02 | 0.27 | 0.07 | T | 0.75 | 0.41 |
| | | | | | | H | 2.87 | 0.13 |
| | | | | | | TxH | 1.90 | 0.21 |
| Extracellular space | | 0.27 | 0.09 | 0.03 | 0.03 | T | 0.95 | 0.36 |
| | | | | | | H | 4.35 | 0.07 |
| | | | | | | TxH | 0.29 | 0.61 |
| Proximity to dark cells | | 0.16 | 0.00 | 0.05 | 0.00 | T | 0.00 | 0.97 |
| | | | | | | H | 1.75 | 0.22 |
| | | | | | | TxH | 0.00 | 0.97 |
| Intracellular organelles | | | | | | | | |
| ER/Golgi | | 7.79 | 1.63 | 7.85 | 0.13 | T | 1.75 | 0.26 |
| | | | | | | H | 0.00 | 0.95 |
| | | | | | | TxH | 3.66 | 0.13 |
| Dilated ER | | 0.87 | 0.00 | 0.89 | 0.37 | T | 2.24 | 0.17 |
| | | | | | | H | 0.00 | 0.95 |
| | | | | | | TxH | 2.24 | 0.17 |
| Mitochondria/ER | | 0.97 | 0.38 | 1.03 | 0.02 | T | 1.59 | 0.28 |
| | | | | | | H | 0.10 | 0.77 |
| | | | | | | TxH | 4.38 | 0.10 |
| Mitochondria | | 3.41 | 0.00 | 2.78 | 0.00 | T | 0.00 | >0.99 |
| | | | | | | **H** | **17.32** | **0.01** |
| | | | | | | TxH | 0.00 | >0.99 |
| Elongated mitochondria | #/cell | 0.33 | 0.04 | 0.46 | 0.17 | T | 3.18 | 0.15 |
| | | | | | | H | 1.94 | 0.24 |
| | | | | | | TxH | 1.94 | 0.24 |
| Dystrophic mitochondria | | 0.08 | 0.00 | 0.32 | 0.09 | T | 0.75 | 0.41 |
| | | | | | | H | 5.59 | 0.05 |
| | | | | | | TxH | 0.78 | 0.40 |
| Primary lysosome | | 0.37 | 0.11 | 0.22 | 0.17 | T | 0.12 | 0.74 |
| | | | | | | H | 3.45 | 0.14 |
| | | | | | | **TxH** | **12.72** | **0.02** |
| Secondary lysosome | | 0.22 | 0.06 | 0.18 | 0.08 | T | 0.05 | 0.83 |
| | | | | | | H | 0.64 | 0.47 |
| | | | | | | **TxH** | **8.64** | **0.04** |
| Tertiary lysosome | | 0.31 | 0.11 | 0.41 | 0.05 | T | 4.35 | 0.07 |
| | | | | | | H | 1.55 | 0.25 |
| | | | | | | TxH | 0.60 | 0.46 |
| Empty phagosome | | 0.34 | 0.07 | 0.53 | 0.01 | T | 0.80 | 0.40 |

| | | | | | | |
|---|---|---|---|---|---|---|
| | | | | | H | 4.26 | 0.07 |
| | | | | | TxH | 0.46 | 0.52 |
| Filled phagosome | 0.06 | 0.01 | 0.10 | 0.06 | T | 0.64 | 0.47 |
| | | | | | H | 0.36 | 0.58 |
| | | | | | TxH | 1.88 | 0.24 |
| Autophagosome | 0.04 | 0.01 | 0.04 | 0.04 | T | 0.36 | 0.56 |
| | | | | | H | 0.00 | >0.99 |
| | | | | | TxH | 1.46 | 0.26 |

| Šídák's multiple comparisons test | | | | | | | | | |
|---|---|---|---|---|---|---|---|---|---|
| | | 1 h *vs* 24 h | | | | Contra- *vs* Ipsi- | | | |
| Parameters | H | p | t | df | T | T | p | t | df |
| BBB contacts | | | | | | | | | |
| Basement membrane | C | 0.36 | 1.39 | 8 | 1.00 | 1 | 0.13 | 2.52 | 8 |
| | I | 0.38 | 1.35 | | 24.00 | 24 | 0.14 | 2.45 | |
| Intracellular organelles | | | | | | | | | |
| Mitochondria | C | >0.99 | 0.00 | 8 | 1.00 | 1 | 0.08 | 2.94 | 4 |
| | I | >0.99 | 0.00 | | 24.00 | 24 | 0.08 | 2.94 | |
| Dystrophic mitochondria | C | 1.00 | 0.01 | | 1.00 | 1 | 0.55 | 1.05 | 8 |
| | I | 0.44 | 1.24 | | 24.00 | 24 | 0.10 | 2.30 | |
| Primary lysosome | C | 0.43 | 1.25 | | 1.00 | 1 | 0.50 | 1.21 | 4 |
| | I | 0.18 | 1.88 | | 24.00 | **24** | **0.04** | **3.83** | |
| Secondary lysosome | C | 0.49 | 1.15 | | 1.00 | 1 | 0.37 | 1.51 | |
| | I | 0.29 | 1.56 | | 24.00 | 24 | 0.11 | 2.65 | |

**Table S9. Microglial ultrastructure relative frequency statistics.**

p values are given after a Fisher's exact test measuring the relative frequency of ultrastructural features per cell across the ipsi- and contra- Lmol and the two timepoints (1 and 24 h). For each data mean and S.E.M of n = 3 animals/timepoint are given. ANOVA: analysis of variance#: number, Lmol: stratum lacunosum moleculare, C or Contra- contralateral, I or Ipsi-: ipsilateral, p: p value and S.E.M. standard error of the mean.

| | | | 1 hour | | 24 hours | | | |
|---|---|---|---|---|---|---|---|---|
| | | | Contra- | Ipsi- | Contra- | Ipsi- | Fisher's exact test | |
| Parameter | unit | | Mean | S.E.M | Mean | S.E.M | Group | p |
| Astrocytic endfeet containing mitochondria | #/cell | Mean | | | | | C1xI1 | |
| | | S.E.M | | | | | C24xI24 | |
| Swollen astrocytic endfeet | | Mean | | | | | C1xI1 | |
| | | S.E.M | | | | | C24xI24 | |
| Extracellular space | Contact / cell | Mean | 0.10 | 0 | 0.20 | 0.03 | C1xI1 | 0.11 |
| | | S.E.M | 0.05 | 0 | 0.06 | 0.03 | **C24xI24** | **0.03** |
| Dystrophic mitochondria | #/cell | Mean | 0.08 | 0.22 | 0.08 | 0.36 | C1xI1 | 0.11 |
| | | S.E.M | 0.04 | 0.07 | 0.04 | 0.08 | **C24xI24** | **0.004** |
| Primary lysosome | | Mean | 0.18 | 0.19 | 0.26 | 0.05 | C1xI1 | >0.99 |
| | | S.E.M | 0.06 | 0.07 | 0.07 | 0.04 | **C24xI24** | **0.02** |
| Secondary lysosome | | Mean | 0.13 | 0.17 | 0.28 | 0.08 | C1xI1 | 0.75 |
| | | S.E.M | 0.05 | 0.06 | 0.07 | 0.05 | **C24xI24** | **0.04** |
| Tertiary lysosome | | Mean | 0.34 | 0.25 | 0.18 | 0.25 | C1xI1 | 0.45 |
| | | S.E.M | 0.08 | 0.07 | 0.06 | 0.07 | C24xI24 | 0.57 |

**Table S10. Microglia ultrastructural nonlinear regression statistics.**

*p* values are given after paired Wilcoxon test of Contra- *vs* Ipsi- nonlinear regression curve fits. These were modeled after the relative distribution of each parameter in the two *LMol* (contra- and ipsi-) nested into the two timepoints (1 and 24 h). For each data, Amplitude, Mean and p value of n = 3 animals/timepoint/hemisphere are given. a.u.: arbitrary unit, *LMol*: *stratum lacunosum moleculare*, #: number, p: p value, Contra-: contra-, Ipsi-: ipsi- and h: hours.

| | | 1 h | | 24 h | |
|---|---|---|---|---|---|
| | | Contra- | Ipsi- | Contra- | Ipsi- |
| | | Synaptic plasticity contacts | | | |
| Synapse containing mitochondria contact (#/cell) | Amplitude | 40.84 | 71.21 | 70.98 | 72.36 |
| | Mean | -0.46 | -3.54 | -0.14 | -0.21 |
| | p | 0.18 | | <0.0001 | |
| Post-synaptic element (#/cell) | Amplitude | 47.54 | 39.60 | 62.38 | 110.50 |
| | Mean | 0.96 | 0.20 | 0.61 | 0.44 |
| | p | <0.0001 | | <0.0001 | |
| Pre-synaptic element (#/cell) | Amplitude | 29.22 | 39.60 | 28.87 | 27.76 |
| | Mean | 2.78 | 0.20 | 1.18 | 0.88 |
| | p | <0.0001 | | <0.0001 | |
| | | Intracellular organelles | | | |
| ER/Golgi (#/cell) | Amplitude | 12.21 | 21.05 | 32.36 | 20.92 |
| | Mean | 7.43 | 4.29 | 3.97 | 4.41 |
| | p | <0.0001 | | <0.0001 | |
| Mitochondria #/cell (#/cell) | Amplitude | 15.50 | 28.00 | 29.44 | 33.34 |
| | Mean | 0.11 | -4.58 | 1.58 | -0.12 |
| | p | <0.0001 | | <0.0001 | |
| Elongated mitochondria (#/cell) | Amplitude | 89.95 | 56.09 | 65.89 | 106.40 |
| | Mean | -0.89 | -0.15 | -0.25 | -0.90 |
| | p | <0.0001 | | <0.0001 | |

**Table S11. Microglial ultrastructural principal component analysis.**

For each PC, eigenvalue, proportion of variance, cumulative proportion of variance and loadings for each morphology parameter of n = 21 cells, N = 3 animals/timepoint/hemisphere are given. PC: principal components

| PC summary | PC1 | PC2 | PC3 |
|---|---|---|---|
| Eigenvalue | 3.252 | 3.026 | 2.363 |
| Proportion of variance | 12.51% | 11.64% | 9.09% |
| Cumulative proportion of variance | 12.51% | 24.14% | 33.23% |
| Component selection | Selected | Selected | Selected |
| Loadings | | | |
| Astrocytic endfeet containing mitochondria contact | 0.77 | -0.07 | 0.19 |
| Astrocytic endfeet contact | 0.85 | -0.15 | 0.14 |
| Swollen astrocytic endfeet contact | 0.62 | -0.15 | -0.14 |
| Basement membrane contact | 0.61 | -0.07 | 0.10 |
| Endothelial cell containing mitochondria contact | 0.37 | -0.13 | 0.15 |
| Microglial blood vessel distance | -0.64 | 0.23 | -0.10 |
| Red blood cell contact | 0.00 | -0.07 | -0.52 |
| Synaptic element containing mitochondria contact | -0.38 | 0.09 | 0.61 |
| Post-synaptic element contact | -0.28 | -0.20 | 0.62 |
| Pre-synaptic element contact | -0.15 | -0.22 | 0.58 |
| Myelinated axon contact | -0.23 | -0.02 | -0.20 |
| Degenerating myelin contact | -0.03 | 0.00 | -0.31 |
| Satellite microglia | -0.03 | -0.01 | -0.39 |
| Extracellular digestion | -0.13 | -0.02 | -0.35 |
| Extracellular space contact | -0.02 | -0.28 | -0.17 |
| ER/Golgi | -0.33 | -0.77 | 0.03 |
| Dilated ER | 0.02 | -0.49 | -0.39 |
| Mitochondria | -0.21 | -0.69 | 0.06 |
| Elongated mitochondria | -0.11 | -0.63 | 0.27 |
| Dystrophic mitochondria | -0.11 | -0.45 | -0.39 |
| Primary lysosome | -0.15 | 0.04 | -0.22 |
| Secondary lysosome | -0.14 | 0.01 | 0.05 |
| Tertiary lysosome | 0.07 | -0.66 | 0.06 |
| Empty phagosome | 0.20 | -0.42 | -0.06 |
| Filled phagosome | -0.23 | -0.44 | -0.07 |
| Autophagosome | -0.10 | 0.10 | 0.09 |

**Table S12. Microglial ultrastructural correlation with blood vessel proximity and blood vessel area descriptive statistics.**

p-values are given after Spearman r correlation between blood vessel area and distance and ultrastructural parameters in the ipsi- *LMol*. For each data, r and p of n = 3 animals/timepoint/hemisphere are given. p: p value, r: r coefficient.

| Ultrastructural parameters | Blood vessel distance | | Blood Vessel area | |
|---|---|---|---|---|
| | r | p | r | p |
| Astrocytic endfeet containing mitochondria contact | -0.6213 | 0.0000 | 0.3151 | 0.0001 |
| Astrocytic endfeet contact | -0.7330 | 0.0000 | 0.3624 | 0.0000 |
| Swollen astrocytic endfeet contact | -0.3126 | 0.0019 | 0.3072 | 0.0001 |
| Basement membrane contact | -0.4722 | 0.0000 | 0.1793 | 0.0276 |
| Endothelial cell containing mitochondria contact | -0.2487 | 0.0146 | 0.1800 | 0.0270 |
| Microglial blood vessel distance | 1.0000 | 0.0000 | -0.0669 | 0.5173 |
| Vessel area | -0.0669 | 0.5173 | 1.0000 | 0.0000 |
| Red blood cells contacts | -0.0662 | 0.5216 | 0.0921 | 0.2606 |
| Post-synaptic contact | 0.0774 | 0.4533 | -0.0807 | 0.3246 |
| Pre-synaptic contact | 0.0082 | 0.9370 | -0.0458 | 0.5768 |
| Myelinated axon contact | 0.0574 | 0.5787 | -0.0635 | 0.4383 |
| Degenerating myelin contact | 0.0658 | 0.5244 | -0.0221 | 0.7878 |
| Satellite contact | 0.0360 | 0.7274 | -0.1194 | 0.1442 |
| Extracellular digestion contact | 0.1584 | 0.1231 | 0.0541 | 0.5097 |
| Extracellular space contact | -0.0988 | 0.3380 | 0.0268 | 0.7436 |
| ER/Golgi | 0.0286 | 0.7819 | 0.0185 | 0.8215 |
| Dilated ER | -0.0142 | 0.8906 | 0.0062 | 0.9398 |
| Mitochondria | -0.0318 | 0.7585 | 0.0577 | 0.4814 |
| Elongated mitochondria | -0.0336 | 0.7453 | 0.1197 | 0.1432 |
| Dystrophic mitochondria | 0.0254 | 0.8060 | 0.0312 | 0.7034 |
| Mitochondria/ER | -0.0875 | 0.3966 | 0.0328 | 0.6894 |
| Primary lysosome | 0.0117 | 0.9096 | -0.0906 | 0.2686 |
| Secondary lysosome | 0.0657 | 0.5248 | -0.0029 | 0.9722 |
| Secondary lysosome | -0.0512 | 0.6204 | 0.0262 | 0.7492 |
| Empty phagosome | -0.0853 | 0.4088 | 0.0322 | 0.6948 |
| Filled phagosome | 0.0191 | 0.8533 | 0.0655 | 0.4241 |
| Autophagosome | 0.0134 | 0.8970 | -0.0944 | 0.2487 |
